# Identification of genetic bases of male fertility in *Pyricularia oryzae* by Genome Wide Association Study (GWAS)

**DOI:** 10.1101/2024.04.25.591152

**Authors:** Alexandre Lassagne, Henri Adreit, Fabienne Malagnac, Florian Charriat, Thomas Dumartinet, Hugues Parinello, Anne-Alicia Gonzalez, Didier Tharreau, Elisabeth Fournier

## Abstract

The reproductive system of an organism impacts the emergence and evolution of adaptive variants in response to selective constraints. The understanding of the sexual mode of reproduction in pathogens helps to understand their life history. In filamentous Ascomycete fungi, mating type system and the production of gametes are required to reproduce sexually. In the phytopathogenic Ascomycete *Pyricularia oryzae*, studies of the genetic determinants of sexual reproduction are still limited to the mating type system. This study focuses on identifying the genes involved in male fertility through the production of male gametes known as microconidia. We performed a GWAS analysis coupled with a local score approach on a wild recombinant population of *Pyricularia oryzae* phenotyped for microconidia production. We identified one genomic region significantly associated with the quantity of microconidia produced. This region contained nine candidate genes, some of them annotated with functions associated to sexual reproduction in model fungi such as *Neurospora crassa*, *Podospora anserina* and *Sordaria macrospora*. The most promising candidate gene contains a Jumonji domain. Proteins belonging to the Jumonji family are conserved among Eukaryotes and are known to be involved in chromatin regulation.

## Introduction

The potential of a pathogen population to adapt to its environment is driven by evolutionary forces, among which recombination is key to reassort existing parental alleles and generate new genotypes (McDonald and Linde, 2002). Sexual reproduction is the main mechanism producing recombination at the genome scale. Thus, the generation of new genotypes thanks to sexual reproduction in pathogen organisms likely contributes to its adaptation to the environment, to the host or to the bypassing of host resistance. In addition to recombination, in several pathogen fungal species, sexual reproduction may lead to the production of specific cells with enhanced survival in unfavorable conditions or during dispersion.

In the heterothallic phytopathogenic fungus *Pyricularia oryzae* (syn. *Magnaporthe oryzae*), the causal agent of blast disease on many cultivated crop with a major food interest (wheat, rice, maize…) (Ou, 1985; Islam et al., 2016; Pordel et al., 2021), sexual reproduction has never been observed in the field, whatever the host specificity of the population. However, evidences from biology, genetic and genomic studies of populations isolated from rice suggest that sexual reproduction took place or is still taking place in limited areas of the Himalaya foothills (the putative center of origin of *P. oryzae*; Zeigler, 1998), some populations present in this area exhibiting equilibrated frequencies of both mating types and footprints of recombination (Saleh et al., 2012; Gladieux et al., 2018b; Thierry et al., 2022). These populations belong to a single lineage. The three other genetic lineages that have spread worldwide are clonal (Saleh et al., 2012; Gladieux et al., 2018b; Thierry et al., 2022) and show all genetic and biological characteristics of asexual populations. Whether the loss of sexual reproduction is a cause or a consequence of the spread of these lineages outside the center of origin is still a matter of debate. To understand the evolution of the reproduction mode of rice blast populations, a preliminary step is to better characterize the genetic determinants of fertility.

In filamentous fungi, the ability to reproduce sexually requires two conditions: the production of functional sexual cells, i.e. male and female gametes, and the recognition, meeting and fusion between these gametes. Recognition and encounter of a male gamete (either contained in antheridia, or present as individualized specialized cells called spermatia) with a female gamete (the ascogonium) is the starting point of the fertilization process. This process is governed by hormonal attraction, and is controlled by a single mating-type locus with two alternative versions in heterothallic fungi (Metzenberg and Glass, 1990). These idiomorphs comprise genes encoding proteins with an HMG-box DNA binding motif that regulates the expression of genes involved in the pheromone/receptor machinery. The mating-type system therefore rules the meeting and fusion of gametes (Debuchy et al., 2010). In *P. oryzae*, the two idiomorphs of the *MAT1* locus (*MAT1.1* and *MAT1.2*) correspond to 2 and 3 genes respectively, one of them encoding a protein containing an alpha-box DNA binding domain (Kanamori et al., 2007). Proteins encoded by HMG-box genes are transcription factors able to bind DNA, facilitate nucleoprotein assembly (Giese et al., 1992), and their role in gamete production vary between species (Koopman, 2010). The mating-type system is not required for the production of gametes in *Podospora anserina* (Coppin et al., 1993) or in *Neurospora crassa* (Ferreira et al., 1998). On the contrary to mating type genes, the genetic determinants governing gamete production in Ascomycetes remain poorly characterized. Some genes involved in spermatia production have been identified, such as *SsNsd* in *Sclerotinia sclerotiorum* (Li et al., 2018) or H3K27 histone methyltransferase gene in *P. anserina* (Carlier et al., 2021). These genes usually have pleiotropic effects. In *P. oryzae*, the loss of function of a H3K4 histone methyltransferase is involved in pathogenicity and more generally in gene activation or repression, but its involvement in male fertility was not tested (Pham et al., 2015). In *P. oryzae*, the implication of the mating-type system in the production of male gametes was not assessed, and more generally, the genetic determinants of this production are completely unknown. In this species, specialized crescent-shaped cells called microconidia were recently shown to be the male fertilizing elements (i.e. the spermatia), and male fertility was therefore defined as the ability to produce microconidia (Lassagne et al., 2022).

Two complementary strategies have successfully been used to identify genes involved in fertility in fungi: reverse and forward genetics. Apart from reverse genetics approaches based on transcriptomic analyses (Garg and Jain, 2013; Riaño-Pachón et al., 2021; Strickler et al., 2012), reverse genetics requires that genes governing the trait of interest have been characterized in one or several model species, and therefore assumes that the genetic processes underlying the trait of interest are similar in the model and focal species. Combined with comparative genomics, such candidate genes approaches brought considerable insights in the understanding of sexual reproduction in Ascomycetes (Ellena et al., 2020; Passer et al., 2022). However, such candidate-genes strategies cannot identify other uncharacterized genes potentially involved in the phenotypic trait of interest. Alternatively, forward genetic approaches allow detecting the genetic determinants of a given trait without a priori. Contrary to reverse approaches, forward approaches aim to link genotypes to phenotypes by detecting statistical correlations between genotypic markers and phenotypic observations in recombinant controlled or wild populations. These approaches have long been restricted to Quantitative Trait Loci (QTL) analyses based on progeny of controlled sexual crosses between parents with contrasted phenotypes, including in fungi (Foulongne-Oriol, 2012). Genome Wide Association Studies (GWAS) approaches have later been used. GWAS was initially developed to face the necessity of understanding human genetic diseases with no access to controlled crosses (Hirschhorn and Daly, 2005). It relies on the detection of significant associations between allelic frequencies at polymorphic positions (Single Nucleotide Polymorphisms, SNPs) along the genome and phenotypic status for a given quantitative trait in wild recombinant populations. It overcomes two major limitations of QTL approaches by avoiding the time-consuming production of laboratory-controlled progenies and by giving access to many more generations of recombination. Within the Fungal kingdom, GWAS proved its efficacy in several model and non-model species, notably in understanding the genetic architecture of adaptation to the host (Dumartinet et al., 2022; Plissonneau et al., 2017; Sánchez-Vallet et al., 2018), resistance to fungicides (Mohd-Assaad et al., 2016; Sanglard, 2019; Spanner et al., 2021), or communication between germinating neighbor conidia (Palma-Guerrero et al., 2013). To our knowledge, fertility in fungi has never been investigated using GWAS. Here, we used this approach to determine the genetic determinants of male fertility, specifically of the production of microconidia in *P. oryzae*. In this purpose, we used full genome sequencing of strains from a wild recombining population for which the production of microconidia was quantified.

## Material and Methods

### Strains of *Pyricularia oryzae*

Seventy-one strains of *P. oryzae* from a single population sampled in the locality of Yule in Yunnan Province of China in 2008 and 2009 (Saleh et al., 2012) were characterized in this study. This population, belonging to the recombinant lineage 1 described in Thierry et al. (2022), was shown to be highly recombinant and suspected to be sexually reproducing (Saleh et al., 2012; Thierry et al., 2022).

### DNA extraction, sequencing, mapping and SNP calling

DNA was extracted as described in Ali et al. (2023). Preparation of libraries and Illumina NovaSeq 6000 sequencing was performed at Montpellier GenomiX (MGX), resulting in paired-end reads of 150 bp. For each sequenced strain, genomic reads were mapped on GUY11 PacBio reference genome with masked repeated elements (Bao et al., 2017; BioSample: SAMN06050153). We choose GUY11 genome (assembly size 42.87 Mb, 56 contigs, contig N50=3.28 Mb) as a reference since this strain belongs to the same genetic lineage as the strains studied here. Mappings were performed with bwa mem v.0.7.17 and filtered with samtools v.1.10 (Danecek et al., 2021) for a mapping quality q=20 (Supplementary text 1: Script1). Mapping quality was assessed with samtools v.1.10 (Danecek et al., 2021) and html-formatting of the report was performed with multiqc v.1.9 (Ewels et al., 2016; Supplementary text 1: Script2). SNP calling was performed using bcftools v.1.10.2 (Danecek et al., 2021) for a minimum depth of 10 reads per position and per individual, and a phred-scaled score above 30. SNPs were filtered for a Minimum Allele Frequency above 5%, and the insertions / deletions were removed. Sites with more than 10% of missing data were deleted (Supplementary text 1: Script3).

### Population structure and LD decay

The population structure of the 71 strains was inferred by a Principal Components Analysis (PCA) using the R packages vcfR v.1.12.0 (Knaus and Grünwald, 2016) and adegenet v.2.1.7 (Jombart, 2008). Linkage disequilibrium (LD) decay was assessed with the PopLDdecay software (Zhang et al., 2019), first by calculating pairwise r² between all pairs of SNPs with a maximum distance of 300 kb, then by averaging all values in adjacent windows of 10 bp. Nucleotide diversity (Pi) was calculated with EggLib v. 1.2 (De Mita and Siol, 2012) from the raw vcf file containing both invariant and variable sites filtered for a maximum missing data of 10% and a minimum depth of 10 reads per position and per individual.

### Microconidia production phenotyping

Microconidia were produced following the protocol described in Lassagne et al. (2022). Briefly, young mycelium grown on Rice Flour medium (20 g of rice flour, 15 g of Bacto agar, 2 g of Bacto Yeast Extract in 1 L pure water with 500,000 U of penicillin G after autoclaving for 20 min at 120°C) was put in 40 mL fresh homemade Potato Dextrose Broth (PDB). PDB was prepared with 200 g of sliced organic potatoes boiled during 30 minutes in 800 mL pure water, filtrated through multi-layer gauze, completed with 20 g glucose and replenished to 1 L. Liquid cultures were incubated during 3 days at 25°C then 6 days at 20°C with permanent shaking (150 rpm). After removing the mycelium by filtration on Miracloth filter film (22 µm), preparations were centrifuged at 4,500g for 15 min and microconidia were pelleted and re-suspended in 1 mL sterile distilled water. The number of microconidia per mL was counted twice with a Malassez cell under optic microscope (X40). Two cultures per strain, considered as independent biological replicates, were carried out. The 71 strains were distributed in 9 batches corresponding to different lots of fresh PDB and dates of experiment. To control for a possible batch effect, 11 strains were duplicated in different batches.

### Statistical analysis of phenotypic values

The microconidia production data were transformed in log(p+1) to reach normality of residues and homoscedasticity. An analysis of variance (ANOVA) using a linear model was performed with the lm function implemented in R. The model used was:

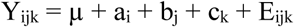

where Y_ijk_ is the logarithm of the production of microconidia, µ is the intercept term, a_i_ the random genotype effect of genotype i, b_j_ the replicate effect in replicate j, c_k_ the effect of batch k and E_ijk_ the residuals term. This model was used to adjust the phenotypic values with the Least Square means (LSmeans) procedure using the lsmeans R package (Lenth, 2016).

### Genome-wide association study and local score approach analyses

GWAS was performed with GAPIT R Software package (version 3) (Wang and Zhang, 2021). As genotypic information, we used the 29 largest scaffolds covering 99% of GUY11 genome. We tested a Mixed Linear Model (MLM) and a Multi-Locus Mixed model (MLMM) (Segura et al., 2012) accounting for genetic relatedness using kinship between strains, estimated using the centered identity-by-step algorithm. The variant component procedure was used to estimate σ^2^_a_ and σ^2^_e_ using restricted maximum likelihood, from the equation

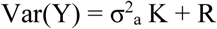

where Var(Y) is the phenotypic variance of the production of microconidia, σ^2^_a_ the genetic variance, K the kinship matrix, and R the residual effect. Under the hypothesis of homogeneous variance, R= Iσ^2^_e_, where I is an identity matrix and σ^2^_e_ is the unknown residual variance. Broad sense heritability defined as the proportion of genetic variance over the total variance was calculated as follows: h^2^ = σ^2^_a_ / (σ^2^_a_ + σ^2^_e_). A p-value of genotype-phenotype association was considered significant when it was smaller than the threshold calculated using the Dunn-Šidák method (Šidák, 1967), calculated as:

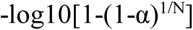

with α the Type I error (here 0.05) and N the number of SNP analyzed.

To get a better resolution on sub-significant peaks of p-values, we used the local score method (Fariello et al., 2017). This method is designed to cumulate local association signals based on p-values found with GWAS method, and is efficient in detecting loci with moderate to small effects (Bonhomme et al., 2019). The aim of the local score method is to identify the genome segments that have a higher density of SNPs with medium to high signal of association, compared with the rest of the genome. The ξ parameter determines the range of p-values contributing to the local score: if p_i_ is the p-value of the i^th^ locus, then the score is taken as X_i_ = −log10(p_i_) − ξ, meaning that only p-values under 10^−ξ^ will contribute positively to the score and p-values above will substract from the signal. P-values distribution was considered as uniform, and we retained ξ=2 and a Type I error risk of α=1 % as local score parameters.

### Analysis of genomic regions associated with microconidia production

Local LD was assessed with the open source software LDBlockShow (Dong et al., 2021) in the genomic regions significantly associated with microconidia production. To precisely identify the candidate genes in these regions, we performed *de novo* gene prediction from the GUY11 assembly using BRAKER1 and AUGUSTUS v3.0.3 software (Hoff et al., 2016), with the same RNA-sequencing data for gene prediction as in Pordel et al. (2021). Gene models of GUY11 (including 2000 bp upstream and downstream of the gene) were compared to their homologs in the 70-15 reference genome (Dean et al., 2005). For the most significant candidate region, to overcome possible errors of annotation, gene models were, when necessary, corrected for start, stop and introns positions with the 70-15 reference genome or *de novo*, so that the coding sequences (CDS) correspond to functional (non-interrupted) protein sequences. For each gene in this most significant candidate region, polymorphic sites in the 71 *P. oryzae* strains were then extracted from the vcf using the coordinates of the corrected gene model in the GUY11 reference genome (Table 1).

**Table 1:** Putative genes in the candidate region associated with microconidia production on GUY11 scaffold 10.

| GUY11 reference genome |  |  |  |  |  | 70-15 reference genome |  |  |  |  |  |  |
| --- | --- | --- | --- | --- | --- | --- | --- | --- | --- | --- | --- | --- |
| Gene ID | Scaffold | Start | Stop | Length (bp) | Strand | Gene ID | Chromosome | Start | Stop | Length (bp) | Strand | Correction<sup>1</sup> |
| Mo_GUY11_054380 | 10 | 927822 | 929318 | 1497 | - | MGG_02051 | 1 | 393105 | 395039 | 1935 | + | No |
| Mo_GUY11_054390 | 10 | 930093 | 931202 | 1110 | + | MGG_02050 | 1 | 391659 | 392768 | 1110 | + | Yes |
| Mo_GUY11_054400 | 10 | 935840 | 938079 | 2240 | + | MGG_02049 | 1 | 384782 | 387021 | 2240 | + | No |
| Mo_GUY11_122940 | 10 | 939051 | 939395 | 345 | + | MGG_16015 | 1 | 383466 | 383690 | 225 | + | No |
| Mo_GUY11_054410 | 10 | 944594 | 946455 | 1862 | - | MGG_02047 | 1 | 376406 | 378267 | 1862 | + | Yes |
| Mo_GUY11_122950 | 10 | 953656 | 954046 | 391 | - | MGG_13692 | 1 | 368872 | 369201 | 330 | + | No |
| Mo_GUY11_122960* | 10 | 958537 | 959340 | 804 | + | MGG_16014 | 1 | 363621 | 364385 | 765 | - | No |
| Mo_GUY11_054420 | 10 | 960874 | 964289 | 3416 | - | MGG_02045 | 1 | 358633 | 362047 | 3415 | + | Yes |
| Mo_GUY11_054430 | 10 | 966804 | 967511 | 708 | + | MGG_16012 | 1 | 355412 | 356119 | 708 | - | Yes |
| Mo_GUY11_054440 | 10 | 967968 | 968239 | 272 | - | NA | NA | NA | NA | NA | NA | No |

<sup>1</sup>: To overcome possible errors of annotation in GUY11 genome, gene models were, when necessary, corrected for start, stop and introns positions, so that the coding sequences (CDS) correspond to functional (non-interrupted) protein sequences.
\*Gene located in a small sub-region where SNP markers were not significantly associated with the phenotype.

To infer haplotype groups, we used Multiple Correspondence Analysis (MCA) implemented in the FactoMineR package (Lê et al., 2008) using the SNP markers significantly associated to the production of microconidia. We performed an ANOVA to test for the effect of the haplotype group on the phenotypic values of the 71 strains, and used Tukey tests for pairwise comparisons of the mean phenotypic value per haplotypic group.

## Results

### Genetic diversity, population structure and recombination within the P. oryzae population

The total number of reads per strain (in million) ranged from 10.9 to 85.9 with a mean of 20.9 and a median of 19.3 (i.e. total sequence per isolate ranging from 1.6 to 12.9 Gb with a mean of 3.1 and a median of 2.9 Gb). Reads were mapped on the GUY11 reference genome with repeated elements masked, and only reads with a mapping quality above 30 were retained. For all strains, at least 75% of the reads mapped on the GUY11 genome, covering more than 80% of it, and leading to a mean mapping depth from 36X to 278X per strain (median 63X; GUY11 genome size of 42.87 Mb according to Bao et al., 2017; Supp. Figure 1). After filtration, 19,331 high-confidence SNPs distributed on the 56 scaffolds of the GUY11 genome were identified among the 71 strains.

The PCA showed no genetic structure among individuals (Supp. Figure 2). Therefore, all strains were kept in further analyses. LD decreased rapidly with physical distance between markers. The fit of a logarithm regression y=a.ln(x) + b (with a = -0.068 and b = 0.9434) to the LD decay resulted in a half-LD decay distance of 3.2 kb (Supp. Figure 3), showing a high recombination rate among the population. Nucleotide diversity within the population, Pi (assessed on 31,834,555 nucleotides including 35,277 polymorphic sites) was estimated at 2×10^-04^ per site. This value agreed with the 2.11×10^-04^ value estimated by Gladieux et al. (2018) for the *P. oryzae* recombinant lineage 1 to which the studied population belongs.

### Phenotypic variability

The strain effect on the production of microconidia was significant, as shown by analysis of variance (F=32.3, P<2×10^-16^, Df=70). Although the production of microconidia seemed to be more variable between replicates for some strains than others (e.g. CH1019; Supp. Figure 4), the effect of the technical replicate was not significant (F=3.43, P=0.067, Df=1). The batch effect, estimated using the 11 strains repeated in different batches, was significant (F=63.3, P<2×10^-16^, Df=8). We corrected for this batch effect using the lsmean method. The corrected value of microconidia production, hereafter expressed in log(1 + number of microconidia per mL), remained highly variable between strains (overall mean, median, and variance: 12.2, 12.0, and 3.5 respectively; Figure 1). Some strains produced high quantities of microconidia, e.g. CH1073, CH1152 and CH1110 (16.2, 16.3 and 17.6, respectively), whereas others were low producers, e.g. CH1093 and CH1126 (8.3 and 8.8, respectively). The broad sense heritability for the production of microconidia was estimated at 0.032.

**Figure 1:**
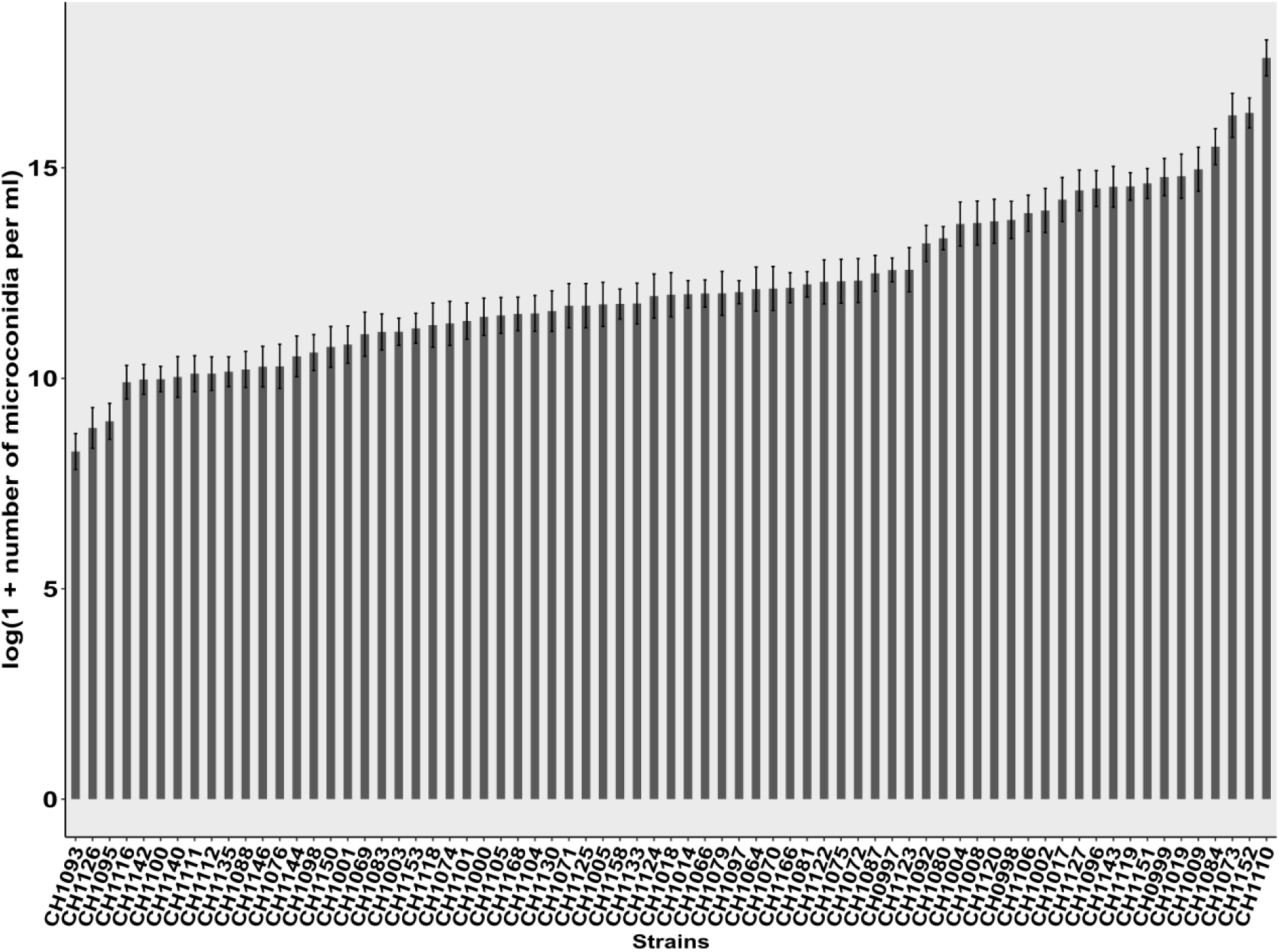
Production of microconidia of 71 strains of *P. oryzae*. The 71 strains were isolated from rice in Yule (Yunnan Province of China). The microconidia production is represented as log(1+ number of microconidia per mL) corrected with the lsmean method. The standard deviation per strain is represented on each barplot.

### Association mapping

GWAS analysis was performed on the 71 individuals using both a Mixed Linear Model (MLM) and Multi-Locus Mixed model (MLMM) that included a kinship matrix. We used the mating-type phenotype (verified biologically by *in vitro* crosses for all strains) as a positive control of GWAS. In the 70-15 reference genome (Dean et al., 2005) the MAT locus is located on chromosome 7 between positions 772,852 and 778,251. The GWAs showed that in the GUY11 reference genome, the mating-type phenotype was significantly associated with a peak on Scaffold 8 between positions 794,254 and 798,440. The alignment of this region against the 70-15 reference genome exactly matched with the position of the MAT locus.

We tested the association between the logarithm of the lsmean corrected values of production of microconidia as phenotypic information [log(1 + number of microconidia per mL)], and the polymorphism of 14,800 SNPs (biallelic, with no missing data) distributed on the 29 largest scaffolds as genotypic information. GWAS with the MLMM identified no SNP marker significantly associated to the microconidia production when the Dunn-Šidák p-value threshold (here: -log[1-(1-0.05)^1/14000^] = 5.43) was considered (Figure 2 upper panel). However, a region of sub-significant p-values was observed on scaffold 10. Within this sub-significant region, the position with the highest -log10(p-value), equal to 3.83, was located at 927,900 bp. This region became significant when the local score approach was applied. (Figure 2 lower panel), and was in fact composed of two very close significant peaks, from position 924,790 to position 955,991 and from position 959,693 to position 967,996, respectively (Figures 3). The non-significant sub-region separating the two peaks started at position 956,006 and ended at position 959,601. In the first peak, the SNP marker with the highest local score (19.3) was located at position 931,007. In the second peak, the highest value was 15.1 at the position 960,405. The highest local scores were 4.6 and 3.6 greater than the local score threshold of 4.2, respectively. The two peaks could be considered as one genomic region of 51.06 kb starting from position 917,291 and ending at position 968,350. This region encompassed 166 SNP markers including 96 SNP markers significantly associated to microconidia production (Figure 3). GWAS with MLM and subsequent local score method leaded to same results as with MLMM (Supp. Figure 5).

**Figure 2:**
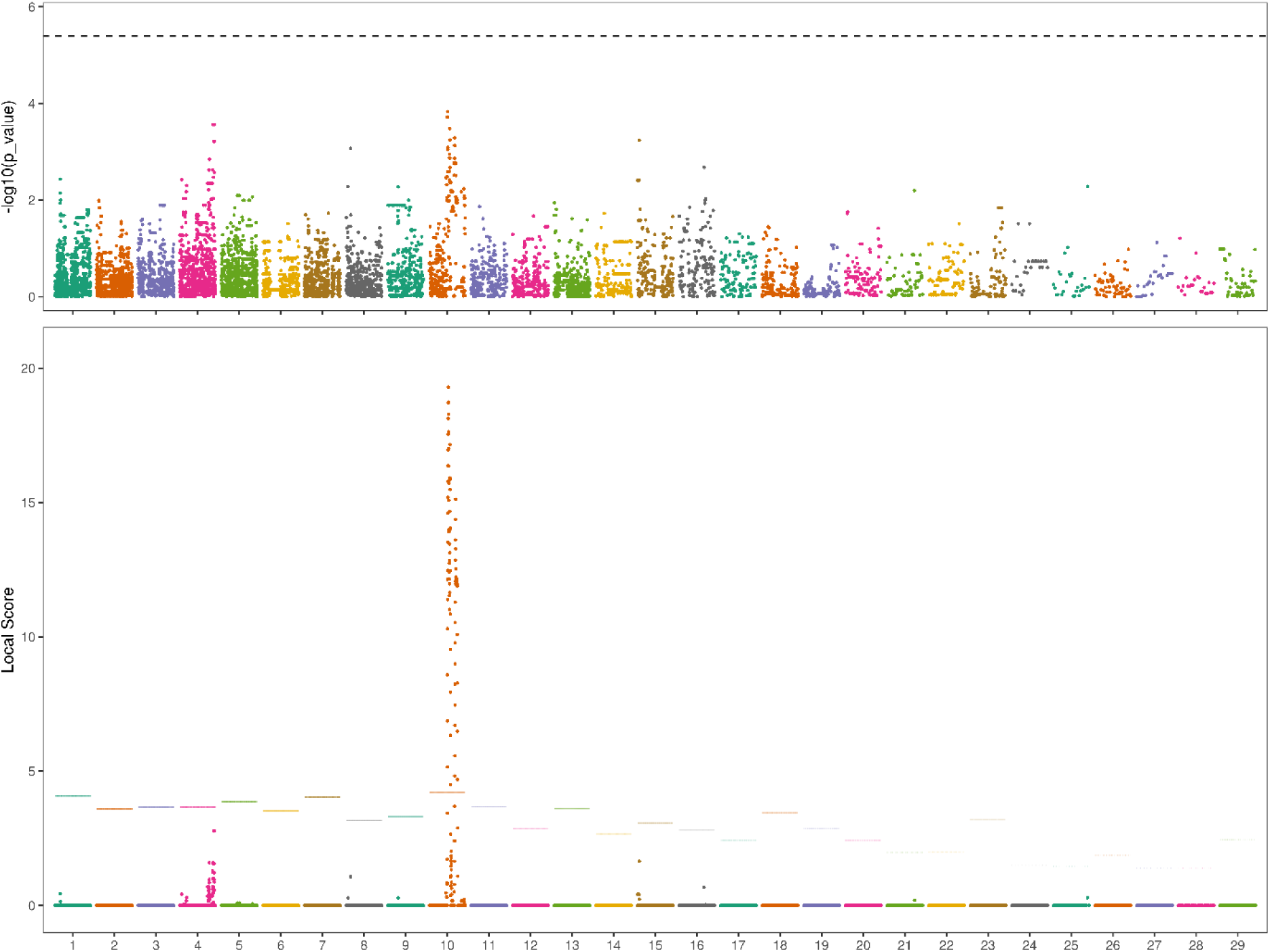
Genome-wide association mapping for microconidia production in *P. oryzae*. Upper panel: Manhattan plot of GWAS showing association p-values for each SNP marker along the 29 largest scaffolds covering 99% of GUY11 genome (one color per scaffold). P-values (vertical axis) are expressed in log. GWAS was performed with a Multi-Locus Mixed model including kinship matrix. The grey dashed line indicates the significance threshold after Dunn-Šidák correction with an α risk of 0.05 (-log[1-(1-0.05)^1/14,000^] = 5.43). Lower panel: Manhattan plot showing SNP marker local score values along the 29 largest scaffolds of GUY11. The local score values were calculated with ξ =2. The horizontal dashed line corresponds to the scaffold-wide local score threshold with an α risk of 0.01.

**Figure 3:**
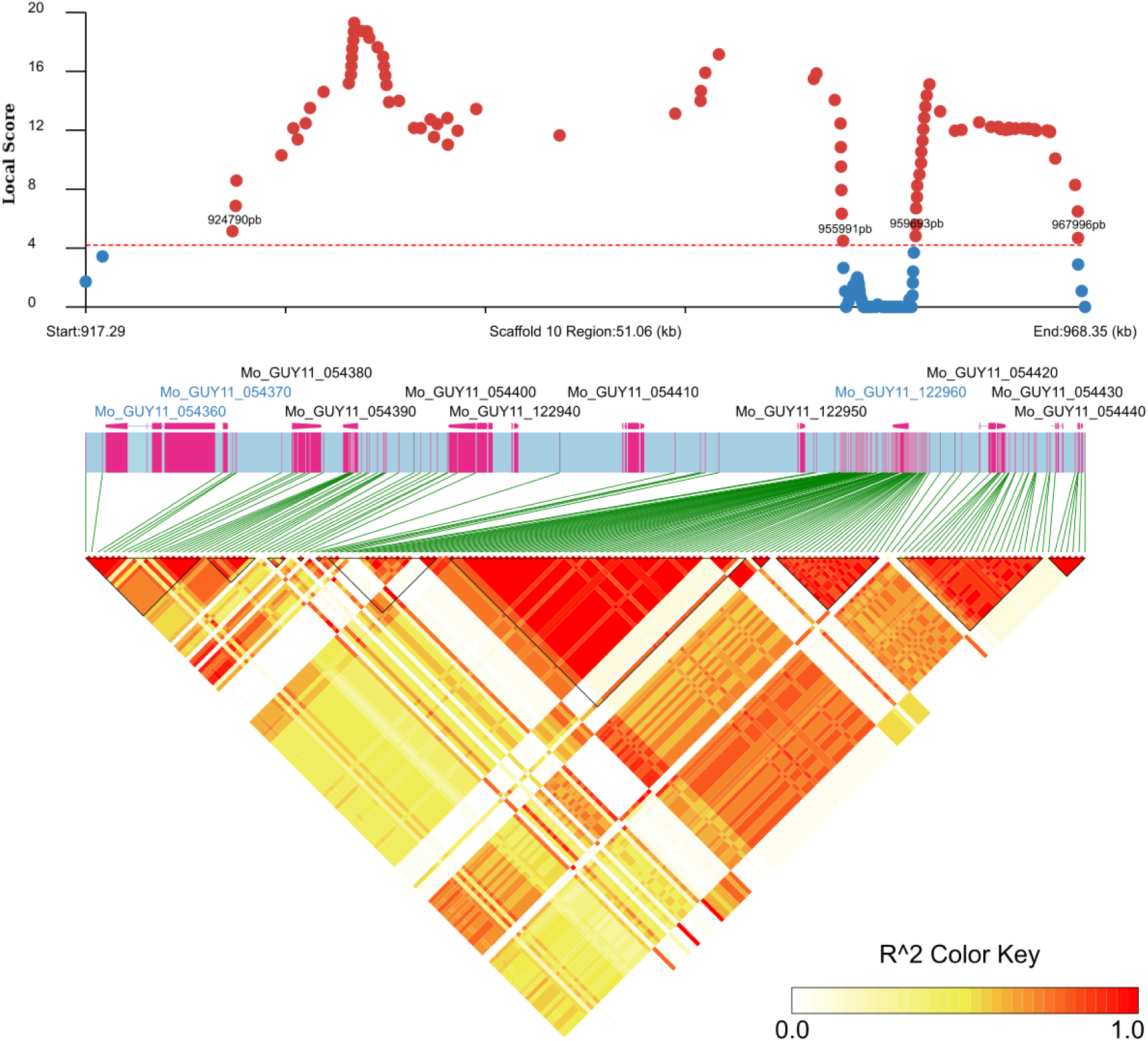
LD blocks contained in the associated region on scaffold 10 determined with the local score method. Upper panel: Manhattan plot of local score analysis on scaffold 10 (red dashed line: scaffold-wide local score threshold = 4.20). Significantly (respectively: non-significantly) associated SNP markers are indicated by red (respectively: blue) dots. Medium panel: Names and structural models of genes models located between the start (917.29 kb) and the end (968.35 kb) of the analyzed region on Scaffold 10. Names in black (respectively: blue) indicate genes located within the significant associated peaks (respectively: outside). Lower panel: LD blocks contained in the 51.06 kb region spanning 166 SNP markers.

The clustering analysis of multilocus genotypes based on the 96 significant SNPs markers defined three haplotypic groups (Figure 4A). The production of microconidia was significantly correlated with the assignation of strains to these haplotypic groups (ANOVA: F=5.35, P = 0.006, Df=2). Haplotypic group 1 produced significantly more microconidia than haplotypic group 3 (Tukey test: P = 0.005), whereas the production of microconidia by strains assigned to the haplotypic group 2 was not significantly different from the two other groups (Tukey tests: 1 vs 2, P = 0.44; 2 vs 3, P = 0.79; Figure 4B). These three haplotypic groups could be considered as three genotypes modulating microconidia production.

**Figure 4:**
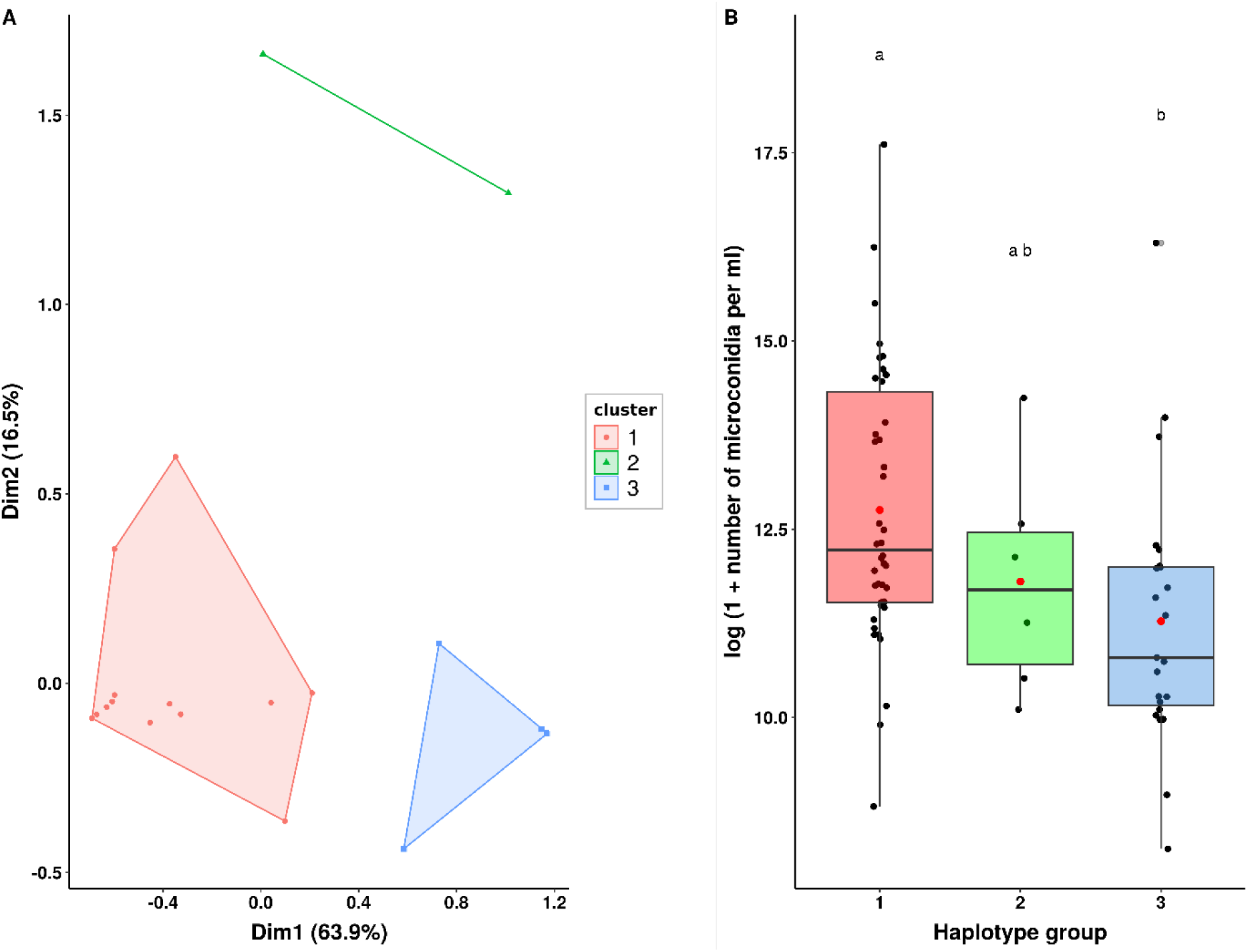
Clustering of strains based on multi-locus genotypes within the significantly associated region on scaffold 10. A: Multiple Correspondence Analysis performed on the 96 SNPs significantly associated to microconidia production, identifying three haplotypic groups (clusters 1, 2, 3). B: Boxplot of phenotypic values of strains assigned to each haplotypic group colored similarly to panel A (mean and median value for each group are shown by red point and horizontal black line, respectively).

### Candidate genes

In GUY11, nine genes are predicted in the genomic region of scaffold 10 located between the first and the last significant SNP markers (*ie* from 924,790 to 967,996; Figure 3). The corrected gene models for each candidate gene in both reference genomes are summed up in Table 1. The markers with the highest local score values were situated in the gene Mo_GUY11_054390 and close to the gene Mo_GUY11_054420 for the first and second peaks respectively.

Among the nine predicted genes (whose sequences in the GUY11 genome are provided in Supplementary text 2), only one had a predicted function in the 70-15 genome (Mo_GUY_054400 / MGG_02049, encoding an Interferon-induced GTP-binding protein Mx2; Table 2). PFAM domains were detected in five other genes: a protein Kinase domain in Mo_GUY_054380 (MGG_02051), a GH43 Pc3Gal43A-like domain in Mo_GUY_054390 (MGG_02050), an oxidoreductase domain in Mo_GUY11_122940 (MGG16015), a WH2 domain in Mo_GUY_054410 (MGG_02047), and a JmjC domain in Mo_GUY_054420 (MGG_02045).

**Table 2:** Annotation of candidate genes.

| Gene ID | Synonymous /<br>Non-Synonymous<br>mutations | CDS<br>length<br>(pb) | Number of<br>alternative<br>proteic<br>sequences | 70-15<br>homologue | Protein function<br>from 70-15<br>annotation | Predicted domain<br>(Pfam / InterProScan<br>/ Uniprot search) | Putative function of<br>homologue protein in<br>other species | NCBI Domain search |
| --- | --- | --- | --- | --- | --- | --- | --- | --- |
| Mo_GUY_054380 | 2/2 | 1497 | 3 | MGG_02051 | Uncharacterized | CAMK protein kinase | CAMK protein kinase | Serine-threo kinase, similar to protein<br>CELE_K09C6 6 |
| Mo_GUY_054390 | 4/4 | 1110 | 2 | MGG_02050 | Uncharacterized | GH43 Pc3Gal43A-like |  |  |
| Mo_GUY_054400 | 0/1 | 2136 | 2 | MGG_02049 | Interferon-induced GTP-binding protein<br>Mx2 | Dynamin-type guanine<br>nucleotide-binding<br>domain, Interferon-induced GTP-binding<br>protein Mx2 | Putative vacuolar<br>sorting protein VPS1,<br>dynamin-2, Dynamin,<br>GTPase domain,<br>mitochondrial fission<br>& membrane fusion | Proteins of the dynamin family catalyze<br>membrane fission during clathrin-mediated<br>endocytosis |
| Mo_GUY_122940 | 0/0 | 276 | 1 | MGG_16015 | Uncharacterized | Oxidoreductase | FAD-binding/transporter-<br>associated domain-like |  |
| Mo_GUY_054410 | 0/0 | 1737 | 1 | MGG_02047 | Uncharacterized | C2H2-type domain,<br>WH2 domain | Actin-cytoskeleton<br>organisation |  |
| Mo_GUY_122950 | 0/0 | 306 | 1 | MGG_13692 | Uncharacterized |  | Breast carcinoma<br>amplified sequence 2<br>(BCAS2) |  |
| Mo_GUY_122960 | 4/7 | 804 | 3 | MGG_16014 | Uncharacterized | DIOX_N domain-<br>containing protein |  |  |
| Mo_GUY_054420 | 1/9 | 3315 | 9 | MGG_02045 | Uncharacterized | JmJ domain | JmjC domain-<br>containing histone<br>demethylation protein | The JmjC domain belongs to the Cupin<br>superfamily. |
| Mo_GUY_054430 | 0/0 | 609 | 1 | MGG_16012 | Uncharacterized |  |  | Septal ring assembly protein ZapB;<br>provisional |
| Mo_GUY_054440 | 0/0 | 177 | 1 | NA | Uncharacterized | NA |  |  |

Only four genes showed polymorphism among the 71 strains in their CDS regions: Mo_GUY11_054400, Mo_GUY11_054390, Mo_GUY11_054380, Mo_GUY11_054420 (Table 2). Among these four genes, Mo_GUY11_054420, which contained the JmjC domain, was the most polymorphic with 10 mutations. Mo_GUY_054420 was located 400 bp downstream of the SNP with the highest local score of the second peak. The ratio of the number of mutations on the CDS length was 0.0030 whereas the mean of the nine genes was 0.0015. Mo_GUY11_054420 contained nine non-synonymous mutations that lead to nine different protein sequences, whereas Mo_GUY_054380, Mo_GUY11_054400, and Mo_GUY_054390 had 2, 1 and 4 non-synonymous mutations respectively (Table 2). The non-synonymous mutation of Mo_GUY11_054400 was in the dynamin GTPase domain.

The region between the two peaks where SNP markers were not significantly associated with the production of spermatia, contained one predicted gene (Mo_GUY11_122960). This gene, which contains a DIOX_N domain, presented 11 mutations in the 71 strains with a ratio of 0.0137 mutation/bp. Mo_GUY11_122960 contained seven non-synonymous mutations that lead to three different protein sequences.

## Discussion

In this study, we phenotyped 71 strains of a recombinant population of *P. oryzae* from Yule (Yunnan Province, China) for male fertility, that is for the quantity of microconidia produced, and showed that this character segregated in the population and was heritable. The use of GWAS combined to local score approach was successful in detecting significant associations between this trait and one genomic region. This region contains nine predicted genes, one of them being a candidate of particular interest. Such a forward GWAS approach remains scarce in fungi, and to our knowledge, has never been applied for studying fertility traits.

The population chosen for this study showed unambiguous genomic footprints of recombination (as shown by LD decay analysis), which confirmed the results obtained by Thierry et al. (2022) with GBS markers. The values of LD decay and nucleotide diversities were also in agreement with the evaluations by Gladieux et al. (2018) for the rice-attacking lineage 1, to which the Yule population belongs. The high recombination rate, high nucleotidic diversity and lack of population structure observed in the Yule population confirmed that it is adequate for GWAS analysis.

The phenotyping of male gamete production showed that this trait, that we equate to male fertility, segregated in the studied population and was heritable, albeit weakly. The significant batch effect highlighted by the analysis of variance indicated that environmental variation could highly impact the production of microconidia, at least in our experimental conditions. Previous studies on other fungal species showed that the production of male gametes tightly relies on available resources (Debuchy et al., 2010; Wilson et al., 2019). Resource availability could result in the complete lack of microconidia production, and consequently, of sexual reproduction, in peculiar conditions. The potential role of environmental conditions in the emergence and spread of clonal lineages in *P. oryzae* (Gladieux et al., 2018; Saleh et al., 2014; Thierry et al., 2022) remains to be deciphered. The environmental effect also likely contributes to the low heritability of male fertility observed in the studied population.

The low heritability of the studied trait might be one reason why classical GWAS failed to detect any significant association between SNPs and the production of microconidia. In our study, the combination of GWAS with the local score approach allowed to circumvent this limitation, proving that local score is a powerful tool for the detection of the genomic bases of weakly heritable traits, and an efficient alternative to QTL analyses. Indeed, QTL approaches are risky for traits related to fertility, because the necessity to cross parents with extreme phenotypes (i.e. in our case: high- / low-microconidia producers) jeopardizes the success of the cross itself and the obtention of enough progeny. Furthermore, QTL approach provides a lower genomic resolution than GWAS and is restricted to the allelic diversity of the two parents (Borevitz and Nordborg, 2003).

In *P. oryzae*, studies on microconidia production and more globally on male fertility are limited to observation (Chuma et al., 2009) and demonstration of their fertilizing role (Lassagne et al., 2022). In *P. oryzae*, mating type genes are required for the formation of perithecia (Wang et al., 2021). Genetic bases of fertility have been scarcely explored for female fertility through perithecia production and asci formation (Lee et al., 2021; Li et al., 2016). Furthermore, in these studies, the genetic bases involved in fertility have been discovered by reverse genetic thanks to homologous genes already identified in other fungal species (Peraza-Reyes and Malagnac, 2016). Here, combining GWAS with local score allowed the discovery of one genomic. The genomic region significantly associated to microconidia production highlighted by our GWAS analysis is a good candidate for further studies. This region, located on GUY11 scaffold 10, is highly resolutive (96 SNPs), and contains a limited number of coding-genes (9).

In the region of interest, Mo_GUY_054420 was the most promising candidate gene. This gene and the marker most significantly associated to the phenotype were physically close (400bp) and belonged to the same LD block. A more detailed analysis of multilocus genotypes in the LD block showed that Mo_GUY_054420 was located in the region where markers are polymorphic between the two genotypic clusters (cluster 1 and cluster 3) formed by strains that significantly differ for microconidia production. In addition, Mo_GUY_054420 was the most polymorphic gene in the region and showed also a high rate of non-synonymous mutations. Mo_GUY_054420 contains a JmjC domain. Although the gene function remains unknown, the Jumonji family protein present in Eukaryotes has been shown to be involved in chromatin regulation and in many signaling pathways (Takeuchi et al., 2006). In *P. oryzae*, the gene MoJMJ1 (MGG_04878) encodes a histone demethylase containing a Jmjc domain. Deletion of MoJMJ1 reduced mycelial growth and asexual spore production, altered germ-tube formation and suppressed appressorium formation (Huh et al., 2017). In the region of interest, we also found the Mo_GUY_054380 gene whose role deserves to be functionally explored with regards to spermatia production. This gene encodes a putative Ca2+/calmodulin-dependent protein kinase (CAMK), a class of serine/threonine-specific protein kinases that phosphorylates transcription factors. Some of the CAMK responding genes were shown to regulate cell life cycle and cytoskeleton network (Berchtold and Villalobo, 2014). These authors also identified the gene Mo_GUY11_054400 which encodes a putative dynamin GTPase potentially related to cytoskeleton organization. Proteins endowed with a dynamin domain are essential to membrane fusion and fission, from endocytosis to organelle division. They are involved in microtubules organization and clathrin-mediated cell membrane invaginations to form budding vesicles (Antonny et al., 2016), a process that may be at work during formation of spermatia from syncytial hyphae. Importantly, none of the five genes of the interval of interest showing polymorphism in their CDS were previously identified as involved in male gametes differentiation.

This study confirmed that the Yule population, and more generally, population from the recombinant rice-attacking lineage 1, are male and female fertile (Saleh et al., 2014; Gladieux et al., 2018; Thierry et al., 2022). Previous studies also showed that the three other rice-attacking lineages found worldwide were clonal, and that populations from these lineages exhibited low to null levels of female fertility and a single mating type. Thus, these populations were considered to reproduce only asexually. It could be interesting to phenotype populations from these clonal lineages for male fertility to test whether they have completely lost the biological ability to produce male gametes, and therefore, to perform sexual reproduction. In parallel, comparing the polymorphism in the candidate genes controlling male fertility in recombinant populations from lineage 1 and in clonal populations from lineages 2-4 would contribute to a better understanding of the causes, stochastic or adaptive, of the loss of fertility that accompanied the migration of clonal lineages all around the world.

## Acknowledgements

We thank Marie Leys, Pierre Gladieux and Stéphane de Mita for their participation in the sequencing process and helpful support in population genomic analyses (calculation of LD decay and Pi). This study was supported by grants from CIRAD, INRAE, ANR-18-CE20-0016 “MagMax”, and the Marie-Curie “EvolMax” project.

## Data accessibility statement

Raw reads will be made accessible at the European Nucleotide Archive (accession no. PRJEB67445). Scripts used for mapping, SNP calling, population genomics analyses and statistical analyses of phenotypic data, are provided as supplementary material.

