## Supplemental data for "Identification of genetic bases of male fertility in *Pyricularia oryzae* by Genome Wide Association Study (GWAS)"

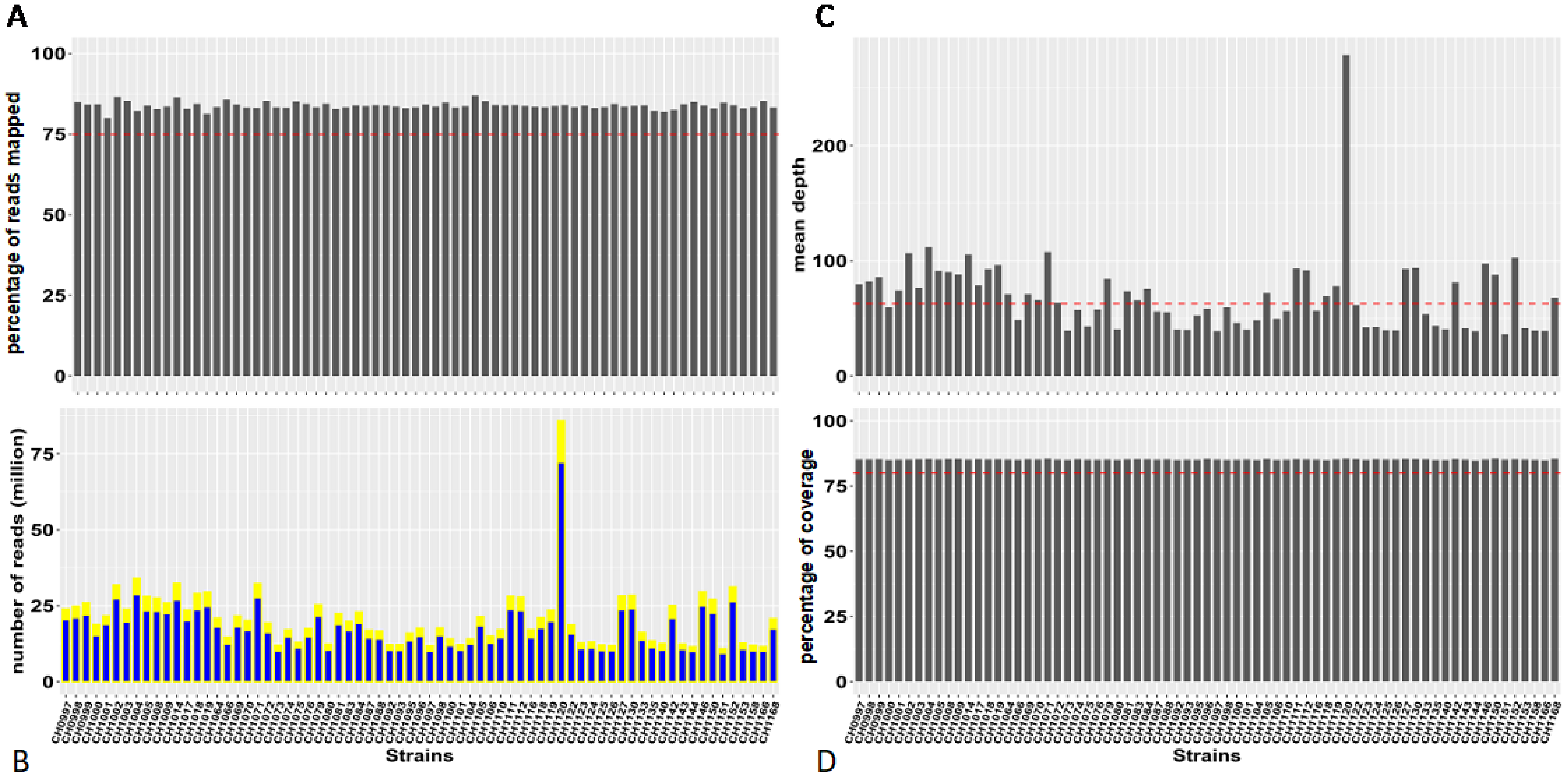


**Supplementary Figure 1: Genome sequencing statistics of 71 *P. oryzae* strains.** A: Percentage of reads mapped on the reference genome GUY11 (red dashed line: 75%). B: Number of reads (in million) produced per strain (yellow + blue) and number of reads (in million) mapped on the reference (blue). C: Mean sequencing depth (red dashed line: median of the 71 values = 63X). D: Percentage of reference genome covered (red dashed: 80% coverage).

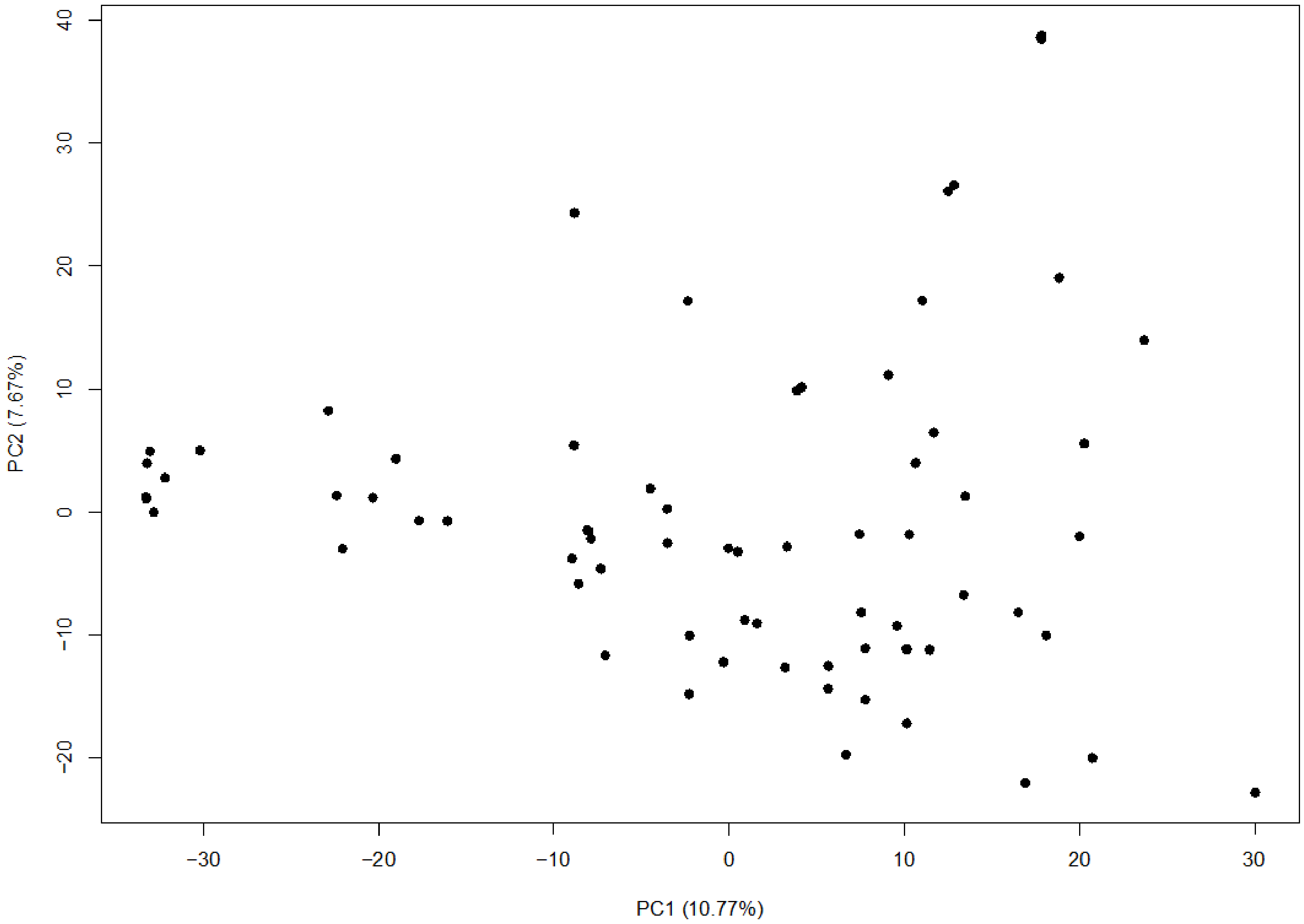


**Supplementary Figure 2: Principal component analysis of the 71 *P. oryzae* strains, based on 14,000 SNPs.**

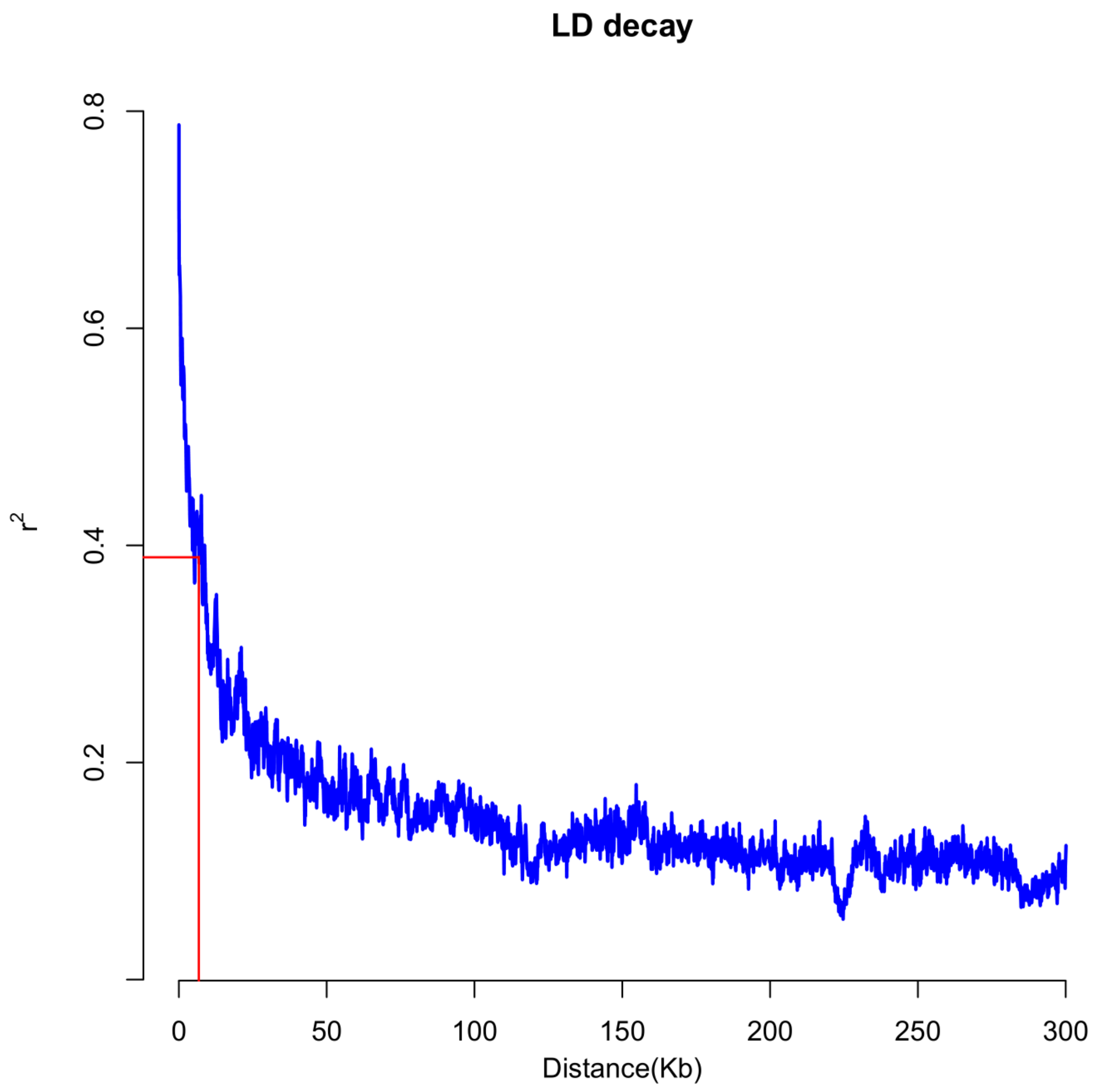


**Supplementary Figure 3: Linkage disequilibrium as a function of physical distance between SNPs among the 71 *P. oryzae* genomes.** The horizontal axis represents the physical distance between markers (in kb). The vertical axis represents the mean value of linkage disequilibrium between equally distant pairs of SNPs averaged in 10 bp adjacent windows. The red line shows the half LD decay, which corresponds to a distance of 3.2 kb and which was calculated after adjusting the data to a logarithm regression y=a.ln(x) + b (with a = -0.068 and b = 0.9434).

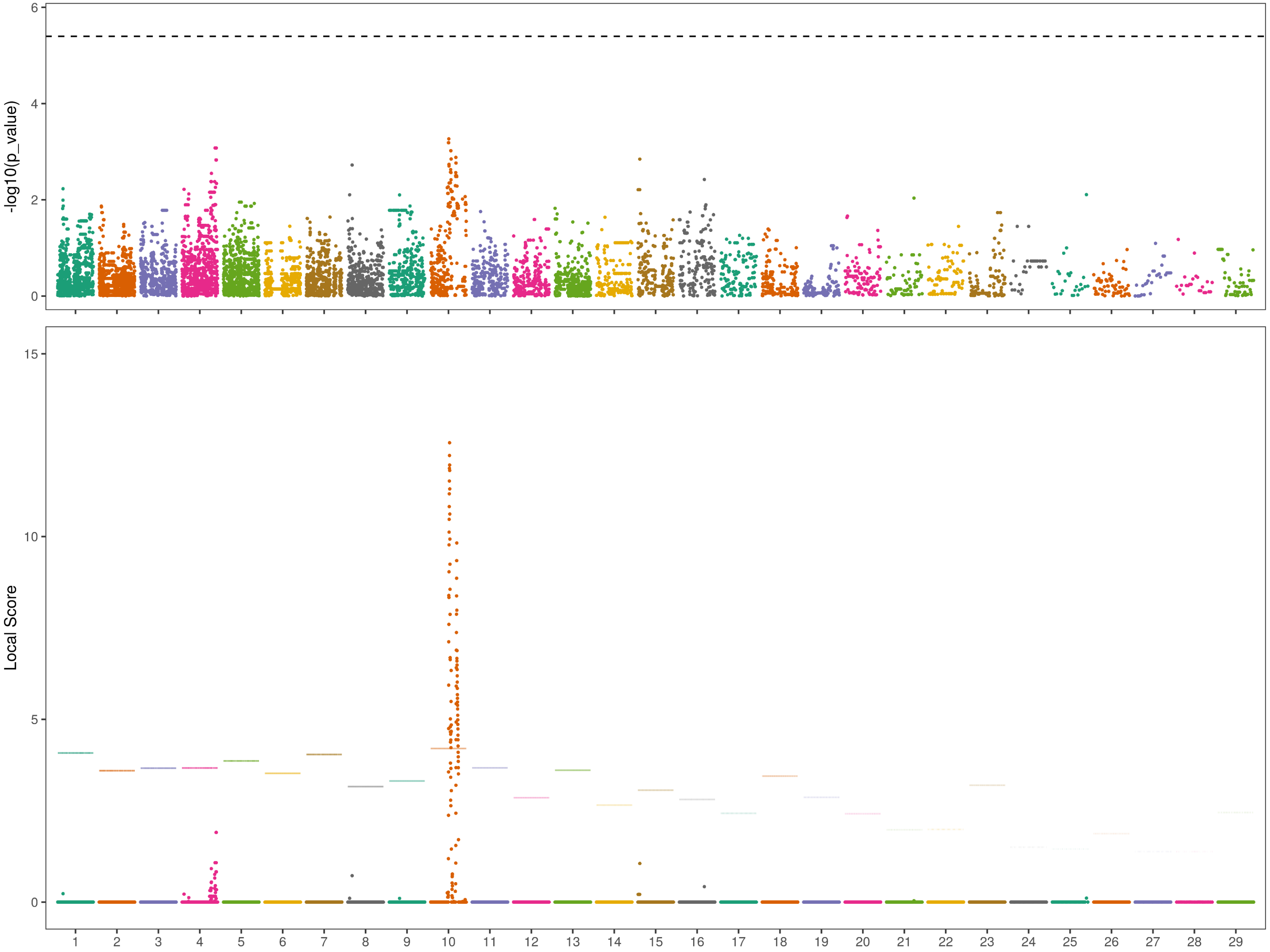


**Supplementary figure 4: Genome-wide association mapping for microconidia production in *P. oryzae*.** Upper panel: Manhattan plot of GWAS showing association p-values for each SNP marker. P-values (vertical axis) are expressed in log. GWAS was performed with a Mixed Linear model including kinship matrix. The grey dashed line indicates the significance threshold after Dunn-Šidák correction with an α risk of 0.05 (-log[1-(1-0.05)1/14,000] = 5.43). Lower panel: Manhattan plot showing SNP marker local score values. The local score values were calculated with ξ =2. The horizontal dashed line corresponds to the scaffold-wide local score threshold with an α risk of 0.01.

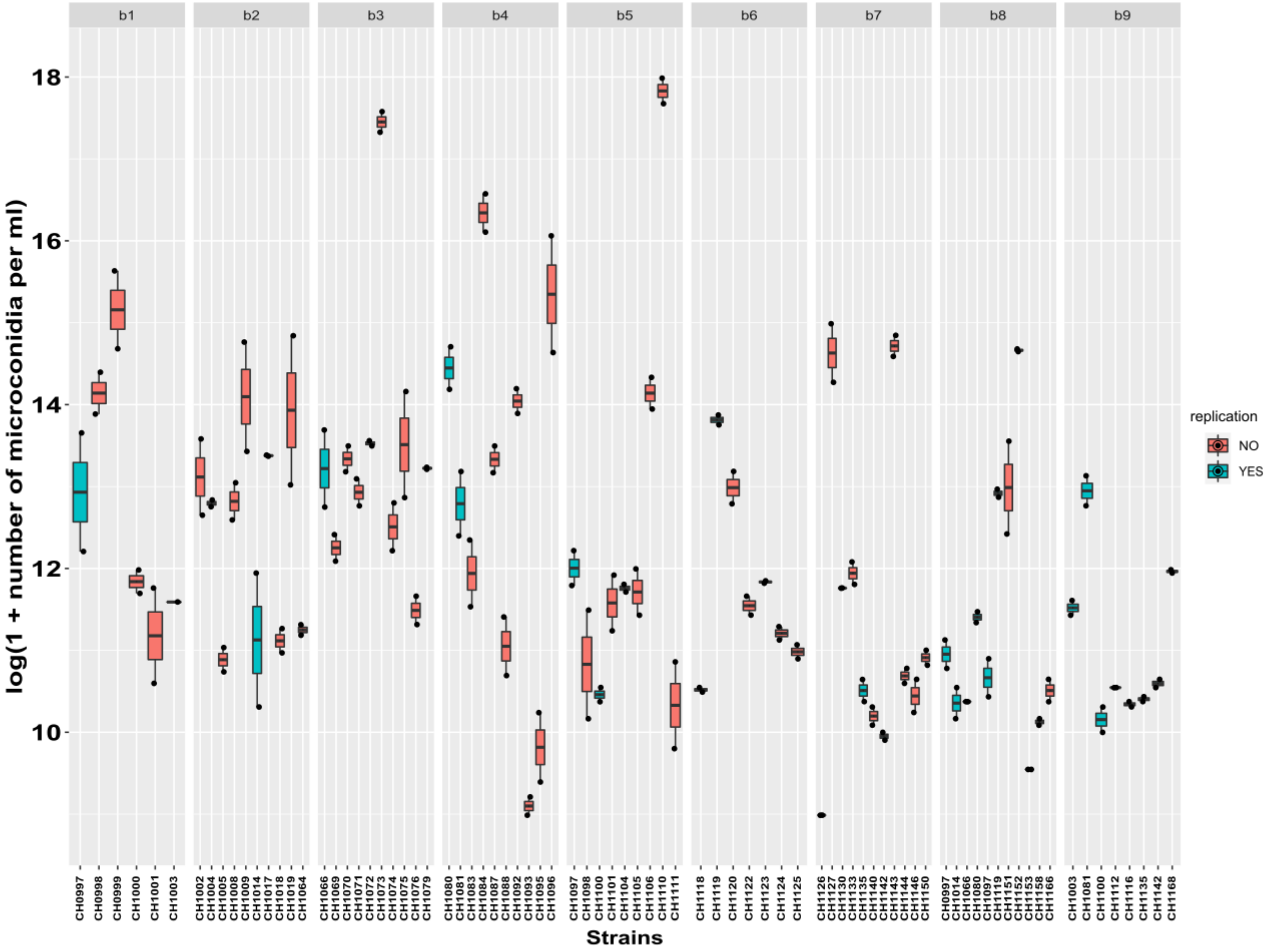


**Supplementary Figure 5: Quantity of microconidia produced by the 71 *P. oryzae* strains.** The nine slots (from b1 to b9) correspond to the different batches of the experiment. The vertical axis is the log(1+ the number of microconidia produced). Blue boxplots correspond to strains replicated in two batches, whereas red boxplots correspond to strains present in only one batch.

### Supplementary text

#### Supplementary text 1

##### Script 1

```
#!/bin/sh
#SBATCH --job-name=mapping
#SBATCH --partition=long
#SBATCH --output=R-%x.%j.out
#SBATCH --error=R-%x.%j.err
#SBATCH --mem-per-cpu=12G

module load bwa/0.7.17
module load samtools/1.10

# Input & Ouput path
REF='/shared/home/alassagne/pyrigenome/ref/GUY11_PacBio_masked.fasta'
PATH_FASTQ='/shared/home/alassagne/pyrigenome/Strains/'
OUTPUT='/shared/home/alassagne/pyrigenome/'


OUTPUT_MAPPING="$OUTPUT/Mapping/"
mkdir $OUTPUT
mkdir $OUTPUT_MAPPING
## Indexing the reference genome
bwa index -a bwtsw /shared/home/alassagne/pyrigenome/ref/GUY11_PacBio_masked.fasta

for fastq in ${PATH_FASTQ}*R1.fastq.gz; do
ID=`basename -s _R1.fastq.gz $fastq`

echo '#!/bin/sh'> $ID.sh
echo 'module load bwa/0.7.17' >> $ID.sh
echo 'module load samtools/1.10'>> $ID.sh

echo "bwa mem $REF ${PATH_FASTQ}${ID}_R1.fastq.gz ${PATH_FASTQ}${ID}_R2.fastq.gz -R
\"@RG\tID:${ID}\tSM:${ID}\tPL:ILLUMINA\" | samtools view -F 0x04 -q 20 -bh | samtools sort
-o ${OUTPUT_MAPPING}${ID}.filt_sort.bam" >> ${ID}.sh
sbatch --export=ALL --partition=long --mem-per-cpu=12G --output=R-%x.%j.out
--error=R-%x.%j.err --job-name=mapping_$ID $ID.sh
done
```

#### Script 2

```
#!/bin/bash
#SBATCH --job-name=mapping_stats
#SBATCH --partition=long
#SBATCH --ntasks-per-node=10
#SBATCH --output=R-%x.%j.out
#SBATCH --error=R-%x.%j.err
#SBATCH --mem-per-cpu=5G

module load samtools/1.10
module load multiqc/1.9

# Input & Ouput path
REF='/shared/home/alassagne/pyrigenome/ref/GUY11_PacBio_masked.fasta'
PATH_FASTQ='/shared/home/alassagne/pyrigenome/Strains/'
OUTPUT='/shared/home/alassagne/pyrigenome/'


mkdir ${OUTPUT}/flagstat
#### Run samtools flagstat and samtools depth on filtered reads
for bam in ${OUTPUT}/Mapping/*.filt_sort.bam; do
ID=`basename -s .filt_sort.bam $bam`
samtools flagstat -@ 10 ${OUTPUT}/Mapping/${ID}.filt_sort.bam > ${OUTPUT}/Mapping/
${ID}_flagstat.txt
samtools depth -a ${OUTPUT}/Mapping/${ID}.filt_sort.bam > ${OUTPUT}/flagstat/
${ID}_depth_a.txt
samtools depth ${OUTPUT}/Mapping/${ID}.filt_sort.bam > ${OUTPUT}/flagstat/${ID}_depth.txt
done
multiqc ${OUTPUT}/flagstat/*.flagstat  -o ${OUTPUT}/flagstat/
#### Estimate mean read depth
# size_ref= total number of position of ref genome size
# sum= sum of depth estimates at each position
# the sum is done only with depth estimated at mapped positions

#### Estimate breath depth = proportion of reads that mapped to the reference genome
# mapped= number of positions of mapped genome

echo -e "ID\tmean_read_DP\tbreath_DP" > ${OUTPUT}/DP_MA.txt
for depth_file in ${OUTPUT}/flagstat/*_depth_a.txt; do
ID=`basename -s _depth_a.txt $depth_file`
mean_read_DP=`awk -v OFS='\t' -v spname="$ID" '{size_ref++; if($3>0) sum+=$3}
END{print spname, sum/size_ref}' ${OUTPUT}/flagstat/${ID}_depth.txt`
breath_DP=`awk -v OFS='\t' -v spname="$ID" '{size_ref++; if($3>0) mapped+=1}
END{print(mapped/size_ref)*100}' ${OUTPUT}/flagstat/${ID}_depth_a.txt`
paste <(printf %s "$mean_read_DP") <(printf %s "$breath_DP")>> ${OUTPUT}/DP_MA.txt
done

#### Estimate the proportion of mapped reads to the reference

echo -e "ID\tMAPPED_reads"> ${OUTPUT}/nbreads_mapped_MA.txt
for flagstat_file in ${OUTPUT}/flagstat/*.flagstat; do
ID=`basename -s .flagstat $flagstat_file`
```

```
awk -v spname="$ID" -F "[+]" 'NR == 5 {print spname, $1}' ${OUTPUT}/flagstat/
${ID}.flagstat >> ${OUTPUT}/nbreads_mapped_MA.txt
done

#in a fastq file, 1read has 4 lines: (nbR1+nbR2)/4 ==> nbR1/2 because nbR1=nbR2
echo -e "ID\tTOTAL_reads"> ${OUTPUT}/nbreads_tot_MA.txt
for fastq in ${PATH_FASTQ}*_R1.fastq.gz; do
ID=${fastq/_R1.fastq.gz/}
nbR=`zcat ${ID}_R1.fastq.gz | wc -l`
TOT=$(($nbR/2))
paste <(printf %s "$ID") <(printf %s "$TOT")>>${OUTPUT}/nbreads_tot_MA.txt
done

paste ${OUTPUT}/nbreads_mapped_MA.txt ${OUTPUT}/nbreads_tot_MA.txt >
${OUTPUT}/prop_mapped_MA.txt
echo -e "ID\tMAPPED_reads\tID\tTOT\tPROPORTION" > ${OUTPUT}/prop_mapped2_MA.txt
awk 'NR>1{PROPORTION = 100*$2/$4 ; print $1"\t"$2"\t"$3"\t"$4"\t"PROPORTION}'
${OUTPUT}/prop_mapped_MA.txt >> ${OUTPUT}/prop_mapped2_MA.txt
```

##### Script 3

```
#!/bin/sh
#SBATCH --job-name=SNPcalling
#SBATCH --partition=long
#SBATCH --output=R-%x.%j.out
#SBATCH --error=R-%x.%j.err
#SBATCH --mem-per-cpu=10G
#SBATCH --cpus-per-task=20
module load samtools/1.10
module load bcftools/1.10.2
module load vcftools/0.1.16

# Input & Ouput path
REF='/shared/home/alassagne/pyrigenome/ref/GUY11_PacBio_masked.fasta'
PATH_FASTQ='/shared/home/alassagne/pyrigenome/Strains/'
OUTPUT='/shared/home/alassagne/pyrigenome/'
OUTPUT_VCF='/shared/home/alassagne/pyrigenome/SNPCAlling_XX.vcf'

bcftools mpileup --threads 20 --annotate "DP,AD,INFO/AD" -f ${REF} ${OUTPUT}/Mapping/
*.filt_sort.bam | bcftools call --threads 20 -m --ploidy 1 -f GQ| bgzip > ${OUTPUT_VCF}
```

#### Supplementary text 2

```
>Mo_GUY11_054380
ATGTCGATACACTGCAAAAGCTCGACGCAGGTGAACTTCGAAATATATTACGATCCCGAGACTGACCAGGTC
ACACTAATAAATGGCCAATCTTCACCCACGCTCGTTCTCAAAAAGGAATCGGGACCCGACAATATTATTGTC
TACAAAGGGACGAGCGCGGAGCTGGAACCTGGCGCCTGGAACCTCACCGCGCCCGATGTTAACCGCACAACC
ACACTTTTGGTTCTCCCACGGCGTTTTCTCCCTGCTAGGGATAATGCTAAAGAATTCGAGCCGGAAAGCAGG
```

```
AAACGATCCGCGTCAGCTACCCTGCCAGCGAAACGTGCGAAGGCCAGCAACACGGTAACATTGGTGCCAAAA
CATACCGTTCCGGTTACCGACTTTAAAATGCGGCAGCTGGAACACCCTCTCGTGTCCCTTAGACCTGGGCAA
ACCCTGAAACTCGTAGGATCGCACCCTCAAAAGGAATATTCCATCACATATGCCCAAAAACTGGCGGAAAAC
AAGTCATCGTTAGTCAGAAAAGTTCAGTTTTCCTTTCCGGGTCATGGAAACGTTGATATGGTTGCCAAAACG
GTACGCAAAGGGATGCGTAACGCTAAAGCTGCTGCCAGGGTCTGGCAGAGCGAAACCAGGATACATAGCAAG
CTGATGGCTACTACGGCTGGCGCCCGTTTCATTGTACAGCTGCTTGGAACAAACGCACGCATCCACACAATA
TACATGGAGGACGTTGAAAGACCGTCACTGGCACACCCTACCTGGCGACAAAGGCCCGAAAACCTGTTTACA
GGCTCACGGGGCGTCGCAGGCCGCATTATGCGTAATATAGCCCTAGCTCTGGATTACGTCCACTCTAAAGGC
ATTACCCATTACGACGTCAAGCCGGGCAACATTCTCTACAACGATACCCGCGGCGCCGTCCTCATCGACTTT
GGTCTCAACAGCGATAGCCCAGATACGGCGACCAATGTTTACGCGGCGGGCACGCCCTGGTATATACCACCC
GAATTTCTTGCGGCATCAAATCCCGGCCGAGGCTTCCCTGGCGACGTCTGGGCACTGGGCGTAGTGCAACTT
TTTCTATTCAAACTGCTTCCGCTCCCGGAAACCACCGCCGAATGGATAATCTCTCATGTCGTTGGTCCGAAA
GGTCCAAACCGCGACAAAGCGAGGGAAACCATGGTTCAATGGCTGAAACGGGTGAAAAATGCCCGGGGATTG
CTTGCCCCAGAGGGCCCTGGATCGGATCCAGACACTTTACTACACGCAGCCGGCGGCCGAAAAGGAAAAGCA
CACCGTAACGGCCAATATAATGCGGTACGAGGCGACGGGGACGGACGAAACCACGACGGTATGGACAAACTT
GTCCACGAGATGACAGAAAGCGACTGCCGCAAACGTATTACAATCCGTAACATCATTGGAGGACTCCGCCCA
GCGGCCAAATACGTGTCGGATGTCGCTCTTCATGATCCTCCCACTACTCCGAATTAA

>Mo_GUY11_054390
ATGGACGGTCAGCGCCCAACTCAACCCGATTTCGACAATGCACCTCAATGGCGCTGGGACTCGTGGAGAGTC
GGACTCGATCCTGAATCGCTTTTCACGATTTTAAGCGAATAATACAACAAGGTCTTTTGCGACATCCAATCC
CCCGAAGCGTTCCATCACGACGTATCTTCAATCGTCCTCCAGGCGAGAAATCTCGAAGATTTCCATCATAGA
TTGTCGAAGCGACGCGAGGAGCGCCTGCGCGAGCTACGAACAGCCTGGACTCGAGTCATACAAAACTTTTAT
GGCTGGCGGCCAATTCGATTTGGCCAAGATGAGGACCTAATATATTCGGTTTTCGGAAAAATGGTTGGGAAC
TTATCCTTCGAGTCTATTATTTGTTATGCCGACGCGCATATACCCGACGACCGCAAATATTCCCCGACCTTG
AGCAACTTGGATCCCGAGGCAAAGCCGCTATCGCCAGACAGCGGACCTGACGCTGGCCCAGACCAGAAGGTG
CTGCAGCGACCCGAAGAGCCCCGGCTGCCGGCGTGGCCGCCAAAGCCAACGACCCGGCCTGCACATGCTTTA
TTAACATCGCCGCCGGGATACTGCAACCGCAAACCGCTCGAAAAGATCGTGCCACCGAGTTTATTGACACCA
GCCGGCGACGATGAGGACGTGACGGTTGAGTCTGCGGCTGCTGGCTCTTCCAAGAGCATACGGAAGCAATCG
AGCACGGAGCAGGGAAAGGAGTGGCAACTCAAGCGGGGACAAAGCCAGGAACTGGAAGCAAAGGAATATAGC
ACTGCACTTAAGCCAGCGGACATCCTGTCCCCTAGCGGCAGCGGCTGCAGCAATCCAAAACGGCAGCAGCAG
AGCATTCAACTGGAGTATGGGCAGGACACTGTTGCAGGCGACGATAAGGACTCACCGTCAAGGACAATGTTT
GCGTCAGCAGTGTTACCACGACGAAAAAGCGCCTATTTGGGGATAACATCGTCACCAACGTTGACCAGCGTT
CGGAGTTGCAAAAAGCGGCCGCGCACACCAGAATCACCCGGCAGTGACGATGGGCCGACCTGCAAAAGGAAC
CGGACGAGAGTCTTCGCCTCGCCGAAGTAA

>Mo_GUY11_054400
ATGCGTAGTGAAGAGCCTAGCGCACCGTCCTCCGTTGTGCGTGGCCTGAACTCGTCCAGGTCGACGGAGCGC
ATCAACCAGATCGATCAAGTAAGAGCCTATGGGATTGGAGATGTTGTCTCTCTACCCCAGCTGGTGGTCTGC
GGCGACCAGTCGGCCGGCAAGAGCTCGGTCTTAGAGAGCGTCACTGGCATCCCGTTCCCCAGAAAGAATGGA
CTTTGTACAAGGTTCCCCACTGAAATCATCCTTCGGCACAACTCCACGGCTACGCGTATTATGGCCACTATT
CACCCTCACGACTCCAGACTGAAGGAAAAGAGAGATGCCCTTCTACAATACCGCCGCGTGCTAGCAGACATG
TCGGAACTCCCTGATGTTATTGAGCAGGTGTCCGCATTGATGGAGATCCGTGGGTATGCGGACCATAGTCTC
GGGAACGCGTTTGCGTCTGATGTCTTGCGGATTGAATTCACGGGCCAGACAAATCTCAATCTGACTATCGTT
GATCTGCCTGGACTCATCTCTGTGGCCAACGAAGAGCAAACAGAAGAAGACATCCAGCTCGTCAAAGACATG
GTTAAAGGCTATGTTCAAAGTTCTCGCACGATTGTGCTAGCAGTGGTGCAGGCCACCAACGACATCGCCAAC
CAAGTCATCATCCAGCTTGCGCGACAGTACGACCCGGAAGGTCAGAGGACGGTGGGGATTATCACCAAGGCG
GACTTGATTAACGAGGGAAGCGAGGCTAGCTTGGCACAGCTTGCGAACAACCAGGCCAACATTAAGCTGAAC
CTTGGCTTTTTTCTTTTGAAAAACCCGAAGCCATCCGAGATTGAAGAGGGTATCAAACCTGAGCAGCGGTCG
CAGAGGGAGCTGAGATTTTTCTCGGAGCCCGTTTGGAAGCAGCATCTCGATATGAGTCGTGTAGGGGCCGAG
AATCTGAAGCTGTTTCTGCAGGACCTCCTGGACGCCCACATCGAGAAGGAAATGCCCCAGGTGGTTGAAGAT
ATTCGGAAGGTCCTGCGCACCACAGAGGAGAGCTTGACGCGACTTGGCCAAGCCCGTCCCACTGTCGGGCAC
ATCCGCATTTTTCTGACGCAAGTTAGCACCAGCTTTGCCCAGCTGGTCCAAGCTGCCTTGGATGGCAACTAT
```

```
CATGGGCCGTACTACGATTTTTTCGATCAGGTTGAAGATTCGCGGTTGCGCGCCGTGGTCCACAAGATCAAT
GGCCAATTCGCTACCGACATCCGTACCCATGGCAAAAAAAGGGTTTTGAGGCCGGCAGATTCAGAGATCAAC
TGGACTCTGGATGAAAATTCAATTAAGGAAAACGACGAGAAGGAGCTTTTCGTGACCCAAAAGGAAATGAAA
AACTGGGTCAAAAAGGTGTATTTGCGCACACGGGGTCGTGAGCTTCCAGGAAACTACAATCATGTCCTGCTG
GCCGAGTTGTTCCTGGAACAATCCTCTCGTTGGCTCCCTATTGCCACAGCCCACGTGGAAAGCATTATCGCG
ATCGTTACAGGGTGGCTGCAGCACGCCACTCGAGTGGTTCTGCCCGAGGATAAACTTCGTGGTGATGTGCTT
TCCATTTGCTCTCAGTGGATTGAGGACGCCGGCAATGACGCTTTGATGGAGCTTGACAAACTTAGAAAAGAC
GAGCAGCGCCAGCCGATTACGTACAACCATTATTACACTGATAACATTCAAAAGTCCCGTCATGGTTTTTTG
CGTGAGGCCGTGGAAGGTGCTATCAAAGAGACAGCAAGCTCGGATTACCATGGCAAACTCCACGTTAGCAAC
ATCCCCATGGATATTGAAAAGTTCCTTGAGGCAATGAAGCGCAAAGTCAACGTCGACATGGACGACCAGGCA
TGTGCTGAGGCCCTGGCGGGGTTGGAAGCTTATTACAAGGTCGCCATGAAGACTTTCGTGGATAATGTCTGT
CGGCAGGTTGTCGAACGTCACATTATTGCGCCCCTTCCCAAGATCTTCAGTCCAGTGACTGTTTCCCGATTC
ACGGACGACGAGCTTCTTCAAATCGGAGCCGAGTCAGAAAAACAAAACCGCAAGCGGGAAGAGCTTAGGAGC
AGGGCAAATAAATTGAGGAGCAGCCTTGAAAACCTCCAGCGACGCTGA

>Mo_GUY11_122940
ATGGACAGCATCAAAAGTTTTGCGGTCGAGAATGGAGCCGACGACGAGTTTCTCTTTCTCAACTACGCCGAC
CTTAGCCAGAACCCTCTGGGAAGCTATGGCGACAAAGACATTGCGTTCATGGAGAAGGTTGCGAGCAAATAC
GACCCGAACGGAGTGTTTCAACGAAATGTGCCTGGAGGATTCAGGTTGTCAATTGCTCGAACAACTGCGTGT
TTGAGGTGA

>Mo_GUY11_054410
ATGGACAACCACCGTTCCCACCAGCCCCTTACAGAAGATAATCTTATCAAACTAAAGGAGCAAATCCCATTA
CTTCCTCGAGAAAGATTGAACAAAGCGAAACGTCGGGTGCATCATCTTATCAGGTGGCTAGAAAATCAATCA
AGAAATGTAGGCATTTTGTTTGCCGTACGTGCACAAATCAACAATGACGACTACGGCTGCCGCGACCACCAT
TACCGAGATCTCGTTTATAAGGGAAATGGCAAACCTGACCTCGATGACCCTAAGTTCTGGGAGGACCAACTT
CGTTCTCTCCAGAAATACAAGCCTGTCGTCAGGATAAAAGAAGCTAAGGGACGCGTCCGTGACGCACTGACT
GGCATTACAAAATTGGGGGGCGAAGGCTTGCAAGTTTTAGAAAATGAGGAGATTTATCTCGGTGAGGGTCAA
GCCCTCCTTTATCGACTCAAAAATGGAGGTTCTACTCCCAACCTCAAAAGCCCATCTTTTTGGGAAGCCAAA
GCCAAACATTTAACATGGCTCTGGAGCACTCTTGAGCGGGAAAAGCATCAACCATCAAAACTTCCGTCACCG
GAGCCACTTTATCAGCCACCGCCCTCCACTGTTTCCTCAGGGCCGCTTTCTGAGATGTTGTCTTGGCCGGCC
CCGGGACCTTCCCTACCCCGGGATCTCCCCAGGAGCCGGCCATGGCGCGAGCCAAGCAGTGAAACCAGCTCC
AACTTCTCCAGCGAGACACCTCCGCCCCACCGTGCCCCGTATCACAAATTTGTCTGGAAAAGCAGGACAGAC
GCTTTCCAGGCGTGGATCGAAACATCCCCAGAAAGTGGTATGCCGTCGCCGTCACCGCCGCCGTCGAACCAT
CTATCGTTCTATGGAGAAGATTTCCCCCTGGGTAGCATGATCATGATCTGGCAAAATCCAGATTTGCAGAAG
TTGGTGGACGCTGAAAAGCGCAACCGGGCTAAATGGTCTGTATACCATTTGCTATCAGGCCTGCAGAACCAT
TGTGGCGAATCAGGTCGCAACGTCGTTGAGGAGGTAGATGACATCTACATCGACGAACGAGGTGCCCACAGA
CGTGGCAATAGAGTAACACCAGTTCCTACGGACGACATTATGGAAACTATCCATCCTCCTGACGAAATGGGA
TTTTCGCGTGTCCAATATGACGCAATGACCCCTGAACAGCGGCAGGAAATTTACAGGAAGAAGGCAAAGATG
CCCAGAGATCAATGGCTTTTAGCCAGGAAGGAGGCAAGAGAGAAGGAGATCGAAATGTACAAAGAGGATCAA
AGAAGGAGAGAGAAGGAGCAAAGGATGATTATTGATGATCTTACGTATTGGGAGGAGAAGGAAGCCTATCTC
GCTGAAAAATACGCAAGCTTGATTGGTGCGGAGCGAAAACCCGGAGACGATGAACTGCCGGACGCTACGCCG
TCAAGAGTTTCAAATAGGAAACGCCGTCACCCAGATGGTGACGCAGATCAAGTAAATTCGCGTGCTGCTACG
CGTCGCAAGTTACAACAGCCAAACTCGACCATCACTCCAGCTTTAACACGACCAACAGCAAGGGCCTCGAAA
CTACGTGAGAGAAGAACAACCACAATCGCCGCTTCAGTAATGGGCAGTTTCTATGACGAAGGTGTGAGGAGA
AGTGCCCGGATTGCGAGGCCGCCAAGGAACCATACTTTGCATGATAGGTCGTCATCCAGTACTAGAGGTACT
AGAAAATGA

>Mo_GUY11_122950
ATGCCACCAATTTCCTCCTCAACCTACACAGACGAGCTTCGATCGCCAGTCTCTCCTGACTACGACCTCCTT
TCCATGCCACCTCCACCATCGACTGGAATCACATCACCCCTAATGGCTGACGCCGGCGAGGCGGAGTTTGTG
GACGGACGGTTGCTGCAAACCCAGCAACAGGGTCTCCACTCCTCGATAGCCAGCAGCGAGCGTGGAAAAACG
CGGCTTCCACCCAATGATTTCTCCAGACATATTCTAAGATCGTCGACCTCATACCCGCAAACCGGTAAACCC
```

```
AACAAAGCTAGCGCCTGA

>Mo_GUY11_122960
ATGTCCTCGACCAGCTGCGTCTCTTTGCCGAAGAGACAGAAATCGTTAAACGGCCGAGTCCCACTAAGGCGG
AAAGGCCGGAAGATATCTTCAGGCAGTACTATGATAGCAAATGGTGACAGAAATACCAAGGCTCTTGTCCCG
GGAGAAGTGTACGCCACGCATTGGGAATACTCCGTTTACTTGGCCATGGTCCTCCCTTGGTCTCCATCGCTT
GCCCAAAACTGTCCGAATATGACTAGGGCTCTGTTTTTTCAACTTATAGCAGCTGACCTGCCCAAAAGGTTA
ATTCCCGAATGCTTTTCGATACTTGACCCGTCTACCTTGAAATGGGCTTCCGGGTACGAAGACGGTGGCGAA
CATGTCAATGAACGCCAATACCCTGTACGATGGTTCGACAAAGAAGGGTTTGTTGGCTGGGTACCAGGTTCG
GAGCTGGTTGGTTTCGACTTTCTCAAGGACTGCGAACAAGAAGAAATACAAGGATTTGGAGACGCCAAAAGA
GAGGACGCAAACACTTGCGGATACAGCAGCTATGCAGAAATGAAAAAGAGGGCAGCCGTTCTTGGAGCGACT
TCAACCCCAATTTCGACGAAGCTCATCTCAGAGATTGCTGCTGACGAGACTCCCATTCCGAAATTTACCTCC
CGCCCAGCAGTTGCTTTAAACGCCAAAGAGCAGTGTGAAATACAGAGCTCCCAACTGACTGGCGAGAGACAC
CAAAAGGAGCTTTCATCATTGGCCAGGGCGGTAAATGCATACATTGATGCCGCAGAAAAGTATCGTCAAGCT
AAGCGAGGATAA

>Mo_GUY_054420
ATGGCAAAACTTAACAGCTTTCTTGATGACGCGCCTGAATCTATTCGCAGTAACAAAGAAGTAATTGCGTGG
CTTTCAGAAAAACTGGAAACGCTAGAATCAGAAAAGAAAATCGAAAGGCTGGAGGCTGAAATCAAGAAGCTC
AAAGGTAGCCGGTCCAACGAAGAGAATCTTCGAAAGCAAGCCAACGAGTGGAAAGCCAAGTATGAAAGACTG
CATAGAGCAGCCACGATTCCAAAGCATGAGCATTCTGAGGATAATGCCAAGCTTATCGAGAAAACCAGACAA
CAGCTGCTCAAGCAGTACGATGAAGAGATTGGGCGTCGGGCCGAGAACGAGCAACGGCAAGCACAGCAGGTT
AGAGAGCAAGCCAAAAAGATCGAGAGATTGCAGGCCGAAAACGCAAAATACAGACAAGTACTCGAAAAGGCC
AAAAAACAAGATTTCCATCCCTCTGCACCAAGTAAAGCGACGTCGGCCAGTGGCAATGTTTACAACGACCTT
ACTGGATATAAAACAAAGTCGTTGACGGTCAGCAACGAAAAGCAAAATGGAAGTCCAAACAATTCCCGCAGA
ACTTGCAGCAGCGTATCCAACGACATCCAGGGTAATGCGCATGCACACGCGGCGGTCAAGCTCGCTATATCA
ACCCTTGATGCCGAATTGCCCGAATCCTACACAAAAAAATTCGATCGATATCGAATCTTTGCAGTTGCCGAGG
CGGCGCCCAGGCCTCAAAGTCCCCCAATTTGAAGGGTCCTGTGAAGTAAGGCTTGGTCAGATGATTGACTGG
GGTACACAATCCTTCCCTAAACGCAAGGCTGAGGTTAGGTACTTTAACGCCGATACATTTATTGGGGCTGGT
GTCTTTTACCAGCTATGTCGCATGGATTCCTGTCGCGTTATTGAGTACGGAAAAAAATTTATCCATCACTGGT
AAATCCCGGAAGCACGCCGACATTACCATTATCCTTTGCCATGTCGCAGCTGGAAACCAGTTGTCGCTCGTA
CAGCCTGACCATTTCGTTGTCGCTATTGTCGACTCCGGAGAAAAGGCTGTGCAACATTTTCAAAAGAAAAGG
GTTCAAGAAGACCGAAAATTGGGGAGCTTGTCCTGCTGGGGAATGGAAAACGCCCTGGTAAACAAGCTAAAA
ACCGATCTCGAACAAGCCTGGAACCTAAATTTGAGTGTTTCTCAGAAAGAGTTTAACTATGCCGGCAATACT
TGTTTACTTCAATGCGTAACATCGGTACGATCATATTTCGGCAAACATACTCCGCAAGAAGGCGACCAATAC
TCTATTCGATTATTTTACCTCGAGCGACTGCTGGAGGAATGGGTCCGTGAAGAAAACGAGTCTGGCACGTCA
AAAACCTCGCAAAACAAACTTCAAAGCCGGACTCCACCAGCAATCATCGCGCTAGCTGAACCATCTTCCCAT
CCCAGCGTATGCGGGGTGGCATATCGGCAGGTTCCTTGCCAGGGTGGCGAATTTGTGCTGCTTCTTCCAAAC
CCTGAACAATATCGGGACCTGAAAGAAAGAAGGAAAGGTAGCCCTTTGCTTTTCACTGTCGCCGAGAAACTC
GGAGCACGAAAGCAGGGCGCATTCCTTATCGAAATCCCCACAGCCTCTCGACCGGAACTTCCCGAACAAAGA
CCTCAGCAAGCACAGTGCACAACCTACGTCCCGGAAAAGCTGGACGGCTTTTGGCGGCTCTTCACTAAAAAA
GAAAAATCAAGCCTTCCTCTTTTACCTGCCAATCCACAGAAGGTTGGCTGTTTGGACGCCGCCACCAAGGCG
TACGAAAACCTCGTTGCGGAATCATTCAAGCAAAAGAAAAGCTTGGAGGGTGTTGCTTATCAGACGGACATC
CCAGCAGAGACCAGCAACGAACGGGATTGTGCTTATTTGCCTCCAGAAAGTCCTATTTGGCCAATCCGAGGC
AACAAGCTACCTCGAACAGGTATACCGGGACTCCATACTCCGACATGCTACCGAGGGACCCCTTTCGCTCCG
TTTGCATGGCATCACGAAGACTTTGAGCTGGGCGCGATCAATTATCTTCATTATGGACAGAAAATATGGTTT
GTCACACACCCAGGGTTCTTTGACAAAGCAACCGCAACGTTTCAAGAAGCTCTAAACATCGACCAAGATCAC
AGCCAATTTCTGCGGCATGAGGCGGTACATGGCGGTGTTCAGTATCTACGGAACAAAGACATACCGACGATA
GGTTTTATGCAAGAAGCCGGGCAAATAGTCGTCGTTTACCCACGCGCCTACCACTCGGGATTTGGTGCGACG
GCGACAGTCGCTGAAGCGGTCAATTATGCCGATCTGCAATACACTCTTCCGCAAGACTACAGGCCGTGTCAC
AAGAAATGCCAGCCGTCAGACCCCATCACCGCTGACTTGGTGTTCCCCGAAAATAATTGTGGGCGGACTGCT
ACCTTACGCTGCAAGCCTAAGACAGTTGCAACTTTGTGCCCCGCGCCGACTTCCCAGGATCTCCGCGAAGGT
GTTCCAAAAAGGAAATACGCACCAGAGCCTAAGAGAAAACCTAGTGAACGGCCTAGATTGAAAGGAAAGAAG
```

```
CACAACGAGGCACAAGACGAACCATCAAACGAGCCCCCAGCGAAACGACAAAAAGACCTACCCCCTCTTGAG
GACATGGCAAACGAGGTCATAAATGCAGAAGAGCGGGCCAAGAGCCACATCATGGCATGTATTGACAACTCG
GATCTCATTTTGCCCAACAAGACTATGGAAAAAAAGGCAGATCAGTATTTAGTCCTTAGCGCACGTTGGGCA
AAGCATGCGCCGTTTAGCAGAATGGTGGCCTTGATATTGGAGACCCACGCAGTTGAGCTGATGTACTATGAA
TCATTTGGCACTTTATTCGACAAGAACAGAGTTTTGCGGCGATTACCAAGCGAAATAGTTGACAAGTCTCTC
GACATGCTGACCGAAAGACCAACCGATTCCGCGTTCAGGACCTGGTTAGGAAGACGCCGTACCTTTTTAAAA
CTTGGCCTGGAATACCTACTGTTTATACCCGTTGCTGGGGAGAAAAGGATGCCTGGCTTTCAAGACTTTGAA
CGGTTAACAGACCATCAGATCCAATCATTGCGCAAACTTATCAAAACCCCTGCCAACGCCGATACCATTGCA
AAAGTCGGCGAAAGATTTCGACGCAGTATTATCGAAGGTGACATCTCGGTTTGGAAAAATATTGGGCAAAAT
GAAATTCGGGATATGAACTTGGAGGAGATATCACGCCGGATTTTTCCGCTCGAAACAGAGCAGGTGGAACGC
TAG

>Mo_GUY11_054430
ATGACCGAAAATATGCCTCCAGGACGTTAAAGTAATACTGAACGCCGGCGAATTAGAGACGACTGTCGCAAA
AGACTGGCTGATCATATCTATGACAGTCTCGGACTCCAAGTCAAACCTGCTGACGTTCGCCTTTTGCCGCGA
GGATCCGATTTTTACAGATGGCGAGTTCTTCCAGGCAACGAGGAAATATGTTCCAAGATATTTTCTAAGAGT
TTGAGTGATCATTCCGTCAACGCATTGCGGCTACTTCGTGAGGAGGTCGGGAAATCGTTTGAAGCGATTGGA
ACGTCAGAATCTCCAAATCTAAATTCTCAGAAGCCGAGTTCGCTTATCAGCCAACTCCAAAATGAAAATGCA
CAACTTTCTGACCAACTGCGTGCTGTAGCGGGAGAACTAGCTACAGAACGTGACCACCGAAGTGATCTGGAA
GAAGAAAATCAAAGTTTGAGGAGCACGCAGAGCTTGTTGGAGAAGAAGCTTCAGGACCAAATAGATATTGCA
GAGAATTTTCGGAGCGTTTTAGCCAGATGTGCTACTGTATTACCTGTTCTCGAAAACATGAGGAAACAAATG
GCGGAATCTTTAGGTGATATTTGTCCAAATTAA

>Mo_GUY11_054440
ATGGGAGCGTTTAAGATTCGAGGCGCGCAGGTGGGGCGAGAAACACAGCCACCGCGAACCAACGCTACTGAG
TGGTCCGCAAGAACGGGCATCCTCATACATCCGACCAATAACCTCTCCAAACACAGTAGTTTTGCCGCGAGC
CACCAGAACCAAACAGCTGCGGTGGGCAGACTCACGAAAGGAGCAAAACAGTCGCATCGCGGGTAG
```
